# SWR1C loss promotes longevity through tRNA-mediated proteostasis

**DOI:** 10.64898/2025.12.05.692690

**Authors:** Ericka Moreno-Mendez, Jimena Meneses-Plascencia, Judith Ulloa-Calzonzin, Ivón Salazar-Martínez, Laura Natalia Balarezo-Cisneros, Nuria Sánchez-Puig, Daniela Delneri, Katarzyna Oktaba, Soledad Funes, Alexander DeLuna

## Abstract

The conserved SWR1C chromatin remodeling complex promotes cellular aging, yet the mechanisms linking its activity to lifespan control remain poorly defined. Although SWR1C shapes chromatin architecture and regulates non-coding RNA expression, how these activities relate to its role in aging remains unclear. Here, we combine genetic and lifespan-epistasis analyses to identify the cellular processes that underlie SWR1C-dependent chronological longevity in *Saccharomyces cerevisiae*. Loss of subunits specifically required for H2A.Z deposition robustly extends longevity, and this effect is functionally linked to cytosolic translation and proteostasis pathways. Lifespan profiling of ncRNA deletions reveals a substantial fraction of aging phenotypes and prevalent genetic interactions with *SWR1*, with tRNAs emerging as key determinants of its long-lived phenotype. The expression of specific tRNA genes is dysregulated in *swr1*Δ cells, and interactions with tyrosine-decoding tRNA genes are linked to ER proteotoxic stress, suggesting that altered tRNA pools affect proteostasis during aging. These findings establish tRNAs as central mediators of SWR1C-associated longevity, revealing a fundamental connection between chromatin remodeling, RNA biology, and proteostasis stress responses in lifespan regulation.

## 1 Introduction

Chromatin remodeling regulates genome accessibility and plays a central role in conserved mechanisms of aging and longevity (Dixon et al. 2015; Pegoraro and Misteli 2009; Swer and Sharma 2021). Across species, epigenetic alterations involving histone composition and chromatin-associated factors are recognized as universal hallmarks of aging (López-Otín et al. 2013). In yeast, histone protein levels decline during replicative aging, and restoring core histone abundance extends lifespan (Dang et al. 2009; Feser et al. 2010). Moreover, the histone variant H2A.Z accumulates with age in mice (Stefanelli et al. 2018) and its overexpression is linked to several pathologies, most notably cancer (Giaimo et al. 2019; Hsu et al. 2018; Svotelis et al. 2010). H2A.Z also accumulates in a sex-dependent manner in Alzheimer’s disease (Creighton et al. 2023), and its removal occurs at distinct genomic sites in young versus aged organisms (Stefanelli et al. 2018). Yet, the downstream cellular mechanisms through which changes in histone abundance, modification, and variant exchange influence aging are not fully understood.

*SWR1*, which codes for a Swi2/Snf2-related adenosine triphosphatase, is the catalytic core of a multisubunit histone-variant exchanger. The SWR1 histone-exchange complex (SWR1C) is conserved from yeast to plants and mammals, where its homologous p400/Tip60 complex integrates histone-exchange and acetyltransferase functions (Hong et al. 2014). SWR1C consists of at least 14 subunits and requires ATP to replace canonical H2A-H2B dimers with H2A.Z-H2B dimers in nucleosomes (Mizuguchi et al. 2004). Intriguingly, only ∼25–30% of genes misregulated in *swr1*Δ and *htz1*Δ (H2A.Z histone variant) mutants overlap in *Saccharomyces cerevisiae* (Mizuguchi et al. 2004; Morillo-Huesca et al. 2010), suggesting both H2A.Z-dependent and H2A.Z-independent roles of SWR1C in gene regulation.

The Chronological Lifespan (CLS) of *S. cerevisiae*, which measures the survival of non-dividing populations, has served as a powerful model for dissecting conserved aging mechanisms (Longo and Fabrizio 2012). Genome-wide CLS studies have revealed hundreds of genetic determinants, highlighting chromatin-remodeling, translation, tRNA modification, and stress responses, among others, as central longevity pathways (Campos et al. 2018; Fabrizio et al. 2010; Garay et al. 2014; Gresham et al. 2011; Matecic et al. 2010). Importantly, the core set of genetic drivers consistently emerging from such screens corresponds to universal regulators of longevity (Cruz-Bonilla et al. 2025). We previously reported that SWR1C impairment extends CLS in *S. cerevisiae* (Garay et al. 2014), and a similar effect of a *swr1*Δ deletion has been observed in *Schizosaccharomyces pombe* (Romila et al. 2021). These findings indicate that loss of SWR1C activity promotes longevity through conserved mechanisms, although the underlying processes remain unknown.

It is known that SWR1C represses pervasive non-coding transcription in *S. cerevisiae*, including ncRNAs transcribed near Pol III loci (Alcid and Tsukiyama 2014), thus emerging as a chromatin regulator that couples histone exchange to RNA polymerase III transcription. Among Pol III products, tRNAs constitute a large and diverse gene family, organized into numerous isoacceptors that decode distinct codons, isodecoders that share the same anticodon but differ elsewhere in sequence, as well as multiple identical gene copies distributed across the genome (Bloom-Ackermann et al. 2014). This remarkable diversity gives rise to dynamic variation in tRNA pools that modulate translation efficiency and fidelity, ultimately affecting elongation kinetics, ribosome pausing, and protein quality control (Komar 2009; Zhang and Ignatova 2011). Recent studies have implicated diverse aspects of tRNA biology in aging and lifespan control across species, including yeast, nematodes, flies, and mammals (Stein et al. 2022; Tyczewska and Grzywacz 2023; Xu et al. 2025; Zhou et al. 2021). Emerging evidence further links Pol III– mediated tRNA expression to proteostatic stress and longevity (Jonak et al. 2024; Malik et al. 2024). Established collections of tRNA and other ncRNA gene deletions have been used to uncover environmentally specific phenotypes (Bloom-Ackermann et al. 2014; Parker et al. 2017, 2018), as well as functional roles of ncRNAs in gene regulation (Balarezo-Cisneros et al. 2021). This resource can now be leveraged to identify novel epistatic links between tRNAs and SWR1C with potential relevance to aging.

Here, we provide a system-level genetic overview of chronological longevity induced by the absence of *SWR1* in *S. cerevisiae*. We show that loss of SWR1C subunits required for H2A.Z incorporation consistently extends lifespan. Using high-resolution aging epistasis screens across more than 600 protein-coding and non-coding RNA genes, we uncovered a functional link between *SWR1* and the cytosolic translation and proteostasis machineries. These CLS-genetic interactions correlate with altered tRNA expression and proteostasis changes, implicating a remodeled tRNA pool in the regulation of proteostasis and lifespan. Together, our results reveal an unrecognized regulatory axis connecting chromatin dynamics, tRNA biology, and protein homeostasis in the control of cellular aging.

## 2 Results

### 2.1 Deletion of genes involved in H2A.Z deposition consistently extends the chronological lifespan of *S. cerevisiae*

Genome-wide surveys of chronological lifespan (CLS) regulators in *S. cerevisiae* reveal that deletion of *SWR1* increases longevity (Garay et al. 2014). More recently, it has been shown that SWR1C impairment increases the CLS of the fission yeast *Schizosaccharomyces pombe*, also by means of large-scale CLS profiling (Romila et al. 2021). To confirm the long-lived effect of the *SWR1* deletion in *S. cerevisiae* under conventional chronological-aging screening conditions, we first used live/dead staining coupled to flow cytometry to measure the survival of *swr1*Δ and WT cells as a function of time in stationary phase. We observed that the half-life of cell populations increased from 3.7 days in the WT to 5.3 days in the *swr1*Δ deletion (**Figure 1A**). To further confirm this phenotype in terms of cell viability, we spotted serial dilutions of the strains at regular time intervals in stationary phase. Compared to the WT strain, *swr1*Δ cells showed higher viability, particularly evident after eight days in stationary phase (**Figure 1B**). These results confirm a reproducible long-lived CLS phenotype for *swr1*Δ in *S. cerevisiae*, independent of the assay used, thereby validating the effect previously identified through large-scale screening.

**Figure 1.**
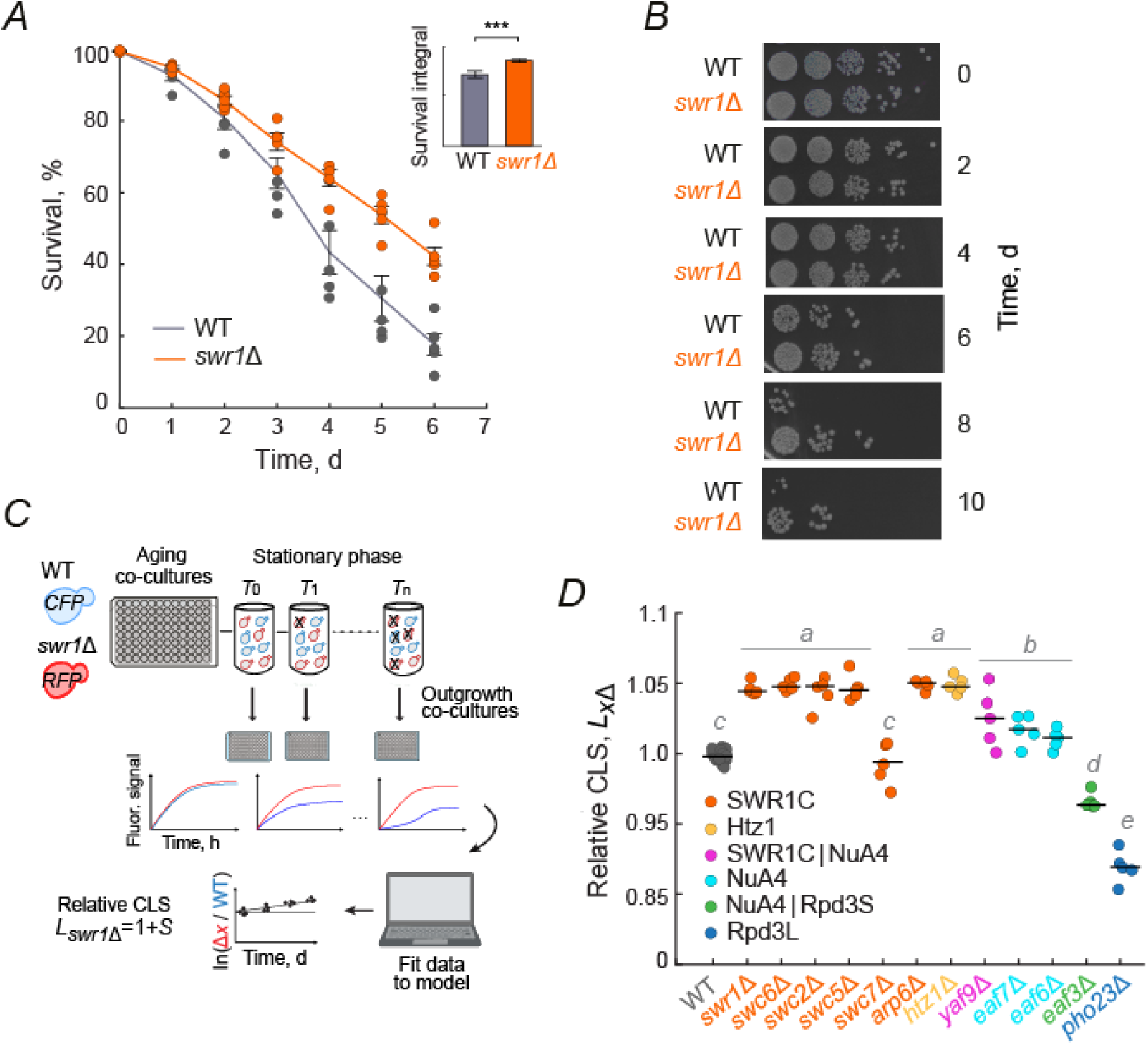
Deletion of genes involved in H2A.Z deposition consistently extends lifespan. (*A*) Percentage of surviving cells inferred by live/dead staining coupled to flow cytometry. Cultures were grown in SC aging medium under full aeration; 100% survival was defined as the population viability 24 h after inoculation (day 0). Error bars are the S.E.M. of four independent experiments. *Inset*: survival integrals (\*\*\**p*=2.2×10^−3^, two-tailed *t*-test). **(*B*)** Viability of WT and *swr1*Δ cells assessed by spotting ten-fold serial dilutions of stationary-phase cultures onto YPD medium after 24 hours of growth, followed by incubation at 30 °C for 48 h before imaging. **(*C*)** High-resolution CLS phenotyping profiling of gene deletions by competitive aging (see Methods). WT and gene-deletion strains were tagged with Cerulean (CFP) and mCherry (RFP), respectively, and grown in co-culture. At stationary phase, samples were collected at designated time points (*T*, days) and outgrowth cultures were monitored at defined intervals (*t*, hours) by measuring OD_600_ and CFP and RFP fluorescence. Survival coefficients, *S_x_*_Δ_, were calculated for each competition experiment using a multiple regression model; the relative CLS was expressed as *L_x_*_Δ_=1+*S_x_*_Δ_. **(*D*)** Relative CLS of single deletion strains of the *HTZ1*-encoded H2A.Z histone variant and components of SWR1, NuA4, Rpd3S, and Rpd3L complexes. Black horizontal lines are the median values from five independent transformation colonies per gene deletion; WT corresponds to the five neutral *ho*Δ transformants, each measured in three technical replicates. Significance was determined by one-way ANOVA followed by Tukey’s multiple comparison test; letters indicate groups that differ significantly (*p*<0.05).

We examined whether the *swr1Δ* long-CLS phenotype is shared by other SWR1C subunits and related chromatin-remodeling complexes. In yeast, SWR1C is composed of at least 14 subunits; part of them are required for the incorporation of the H2A.Z histone into nucleosomes *in vivo* through the ATP-dependent exchange of canonical H2A-H2B into H2A.Z-H2B dimers. Therefore, we first asked whether lack of the *HTZ1* encoded H2A.Z variant had similar CLS impacts, given that only about 25-30% of the regulated genes overlap between the *swr1Δ* and *htz1Δ* deletion strains (Mizuguchi et al. 2004; Morillo-Huesca et al. 2010). To increase the resolution of our assays, we used a previously reported competitive-aging screening approach (Avelar-Rivas et al. 2020) that provides a quantitative measurement of the CLS effects of gene deletions (**Figure 1C**). Deletions of all SWR1C subunits, as well as *htz1Δ*, resulted in similarly extended CLS phenotypes that were statistically indistinguishable from one another and significantly longer than WT (**Figure 1D**). Importantly, Swc7, the only SWR1C subunit whose deletion displayed a WT phenotype, is also not required for Htz1 deposition (Kobor et al. 2004; Lin et al. 2017).

The acetyltransferase NuA4 and SWR1C complexes are physically associated in plants and animals in the homologous TIP60/p400 mammalian complex, which include both histone acetyltransferase (Tip60) and histone exchange (p400/Domino) activities. In this context, pre-existing chromatin acetylation by NuA4 promotes the substitution of H2A with H2A.Z (Altaf et al. 2010). These previous findings suggested that changes in histone acetylation promoted by the NuA4 complex could result in similar effects on yeast CLS as those observed in the absence of SWR1C. To test this hypothesis, we evaluated the CLS in mutants of subunits specific to NuA4 (*eaf3Δ, eaf6Δ*, *eaf7Δ*), some of which are shared with SWR1C (*yaf9Δ* and *bdf2Δ*), and *pho23Δ*, a member of the unrelated Rpd3 histone deacetylase complex. We found that mutants of subunits shared between SWR1C and NuA4, and those directly involved in the acetyltransferase activity of NuA4 (*eaf6Δ* and *eaf7Δ*) also extend yeast longevity, although to different quantitative extents (**Figure 1D**). In contrast, the deletion of *PHO23* led to a decreased lifespan. Taken together, these results show that deletion of genes required for H2A.Z deposition consistently increases lifespan, establishing the conserved SWR1C complex as a key aging factor whose impairment promotes cellular longevity.

Although SWR1C impairment extends lifespan in both *S. cerevisiae* (Garay et al. 2014) and *S. pombe* (Romila et al. 2021), mutant phenotypes often vary widely across individual yeast strains. We therefore tested whether SWR1C loss also promotes longevity across multiple *S. cerevisiae* backgrounds. To this end, we deleted the defining catalytic member of the complex in genetically distant strains that were isolated from different environments and measured their CLS. Specifically, we generated *swr1*Δ transformants in four different strains; L-1374 and Y12 are from wine must in Chile and palm wine in Ivory Coast, respectively, while YJM978 and YJM981 are isolates from vaginitis patients (Liti et al. 2009). By comparing the survival curves with those of the corresponding WT strain in each background in individual cultures, we observed a significant long-lived phenotype of *swr1*Δ in three out of the four strain backgrounds tested (*p*<0.05; **Figure S1**). The corresponding gene deletion in the YMJ981 strain also showed a modest increase in CLS compared to its WT background, but this difference was not significant. Together, these data indicate that SWR1C-dependent longevity is not strain-specific but instead represents a broadly robust phenotype across diverse *S. cerevisiae* genomic backgrounds.

### 2.2 Longevity induced by SWR1C impairment is mediated by translation, proteostasis, and stress-response pathways

Genetic interactions provide a valuable entry point for inferring gene function in the absence of prior knowledge. To investigate the biological processes underlying *swr1*Δ-induced longevity, we used a synthetic genetic array (SGA) approach to generate a double deletion library by combining *swr1Δ* with a curated set of 314 aging-associated genes (**Figure S2**; **Dataset S1**). This gene set includes the major cellular components and biological processes linked to chronological aging, as defined by a recent meta-analysis of genome-wide CLS screens (Cruz-Bonilla et al. 2025). Using our competitive-aging screen, we quantified the effects of each gene deletion in both *SWR1* and *swr1Δ* backgrounds. The CLS measurements were highly reproducible between two independent experiments (*r*=0.93; **Figure S3A**), supporting the robustness of our dataset for high-resolution mapping of genetic interactions underlying SWR1C-mediated lifespan phenotypes.

Using the ensuing quantitative CLS data, we asked which gene-deletion phenotypes were modified by the absence of *SWR1*. Specifically, lifespan-epistasis interactions, ɛ, were defined deviations—either positive or negative—from the null model of no epistasis, where the lifespan of the double mutant equals that of the single deletion *L_xΔswr_*_1*Δ*_=*L_xΔ_* (**Figure 2A**; **Dataset S1**). For example, deletion of *RPS14A*, coding for a small ribosome subunit protein, had a deleterious effect on CLS relative to WT, but this effect was rendered neutral compared to *swr1Δ* in the *rps14aΔswr1Δ* double mutant (ɛ=0.146, positive epistasis). Conversely, deletion of *FAR11*, a gene involved in recovery from cell cycle arrest, had a neutral effect alone but became detrimental in combination with *swr1Δ* (ɛ=‒0.081, negative epistasis). These high-resolution lifespan-epistasis profiles thereby uncover functional associations between the SWR1C and diverse cellular processes that shape aging phenotypes.

**Figure 2.**
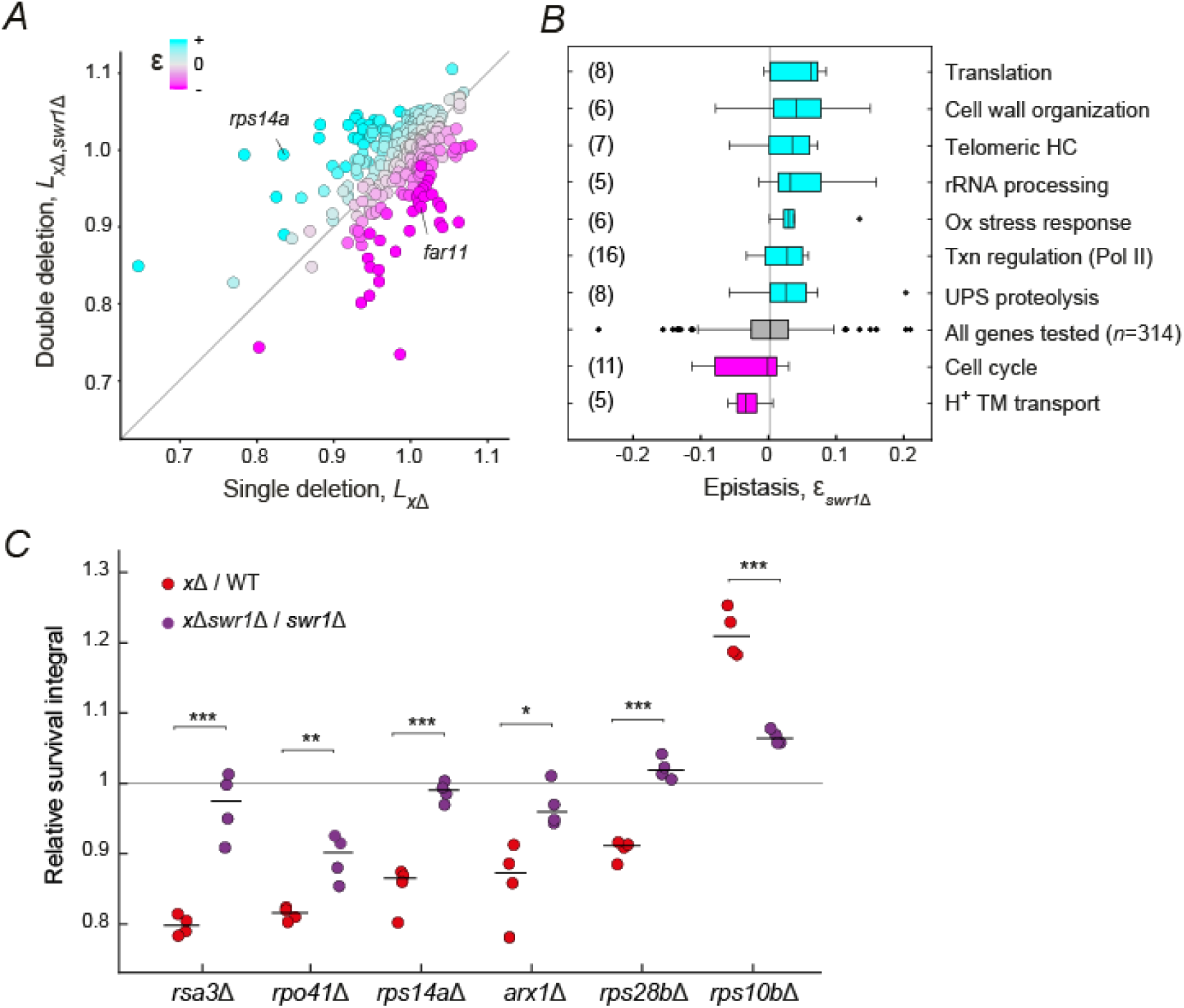
Lifespan epistasis profiling reveals a functional link between *SWR1* and the translation and proteostasis machineries. (*A*) Scatter plot showing the CLS of 314 single mutants, *L_xΔ_*, and corresponding double mutants in competition with *swr1*Δ, *L_x_*_Δ*swr1*_*Δ*. The color scale represents epistatic genetic interaction, defined as ɛ=*L_x_*_Δ*swr1*Δ_‒*L_x_*_Δ_. **(*B*)** Functional analysis of epistatic interactions between *SWR1* and 314 aging-associated genes. Each bar represents the median ɛ value for genes grouped by Gene Ontology (GO) annotation, compared to the global median of the dataset. GO groups of at least five genes with a significant positive (cyan) or negative median ɛ (magenta) are shown in the plot (*p*<0.05, Wilcoxon rank-sum test) (**Table S1**). The gray box plot represents the ɛ values of all screened genes and the corresponding vertical gray line is their median ɛ value. *Abbreviations*: HC, heterochromatin; Ox, oxidative; Txn, transcriptional; UPS, ubiquitin-proteasome system; TM, transmembrane. **(*C*)** Validation of selected epistatic hits. Replicates of single and double mutants were aged in individual cultures, and survival curves were obtained using live/dead staining of stationary-phase cultures at different days. The plot shows the survival integrals (AUC) for single and double mutants, expressed relative to the average AUC of the WT (red) or *swr1*Δ (purple), respectively. Pairwise statistical comparisons between single and double mutants were evaluated using two-tailed *t*-tests (\**p*<0.05, \*\**p*<0.01, \*\*\**p*<0.001).

We examined which groups of genes were associated with positive or negative lifespan-epistasis interactions with *SWR1* (**Figure 2B**; **Table S1**). The results of our functional genetic analysis were consistent with previous gene-expression, ChIP, and genome-wide mutant screening analyses, showing that impaired H2A.Z deposition affects genes involved in cell wall biogenesis, RNA polymerase II transcription, telomeric heterochromatin, and cell cycle regulation (Caydasi et al. 2023; Mizuguchi et al. 2004; Morillo-Huesca et al. 2010; Papamichos-Chronakis et al. 2006; Zhou et al. 2010). However, it was unexpected to find that many of *SWR1*’s genetic interactors in our CLS-based phenotyping screening were genes associated with cytosolic translation, ribosomal RNA processing, ubiquitin-mediated protein degradation, and oxidative stress responses.

To validate these findings, we evaluated six genetic interactions involving genes related to translation and ribosomal maturation by measuring the CLS of single and corresponding *swr1*Δ double mutants using live/dead staining of stationary-phase cell cultures (**Figure 2C**). All short-lived single knockouts tested showed a significant lifespan extension in the *swr1*Δ background, in most cases reverting to a neutral relative phenotype, indicating that the short-CLS phenotype of mutants impaired in cytosolic translation requires the presence of *SWR1*. Conversely, the long-lived phenotype of the *rps10b*Δ mutant was suppressed in the double mutant with *swr1*Δ, suggesting that deletion of both genes extends CLS through a shared mechanism. Together, these results indicate that altered dynamics of protein synthesis contribute to the extended CLS phenotype of *swr1*Δ cells.

### 2.3 Deletion of *SWR1* exacerbates translation stress without altering global translation rates

To further examine the connection between *SWR1* and cytosolic translation, we tested growth sensitivity to cycloheximide and hygromycin B, translation inhibitors that expose defects in strains with compromised translational capacity (Borovinskaya et al. 2008; Schneider-Poetsch et al. 2010). The *swr1Δ* strain displayed marked sensitivity to both drugs, confirming a functional link between SWR1C and cytosolic protein biosynthesis (**Figure 3A**). To evaluate whether SWR1C influences longevity by altering global translation rates, we compared polysome profiles from WT, two SWR1C mutants (*swr1*Δ and *arp6*Δ), and the ribosomal-protein mutant *rps18a*Δ, included as a positive control for impaired translation. Polysome distributions in *swr1Δ* and *arp6Δ* were largely indistinguishable from WT, and the polysome-to-monosome ratio (P/M) remained unchanged, indicating that the global balance between translation initiation and elongation is preserved (**Figure 3B**). In contrast, the *rps18a*Δ control displayed an imbalance in 60S and 40S subunits, consistent with reduced translation initiation and diminished overall translational activity. These findings show that SWR1C loss perturbs translational homeostasis without causing major defects in overall ribosome occupancy on mRNAs, although shifts in the relative synthesis of specific proteins cannot be ruled out.

**Figure 3.**
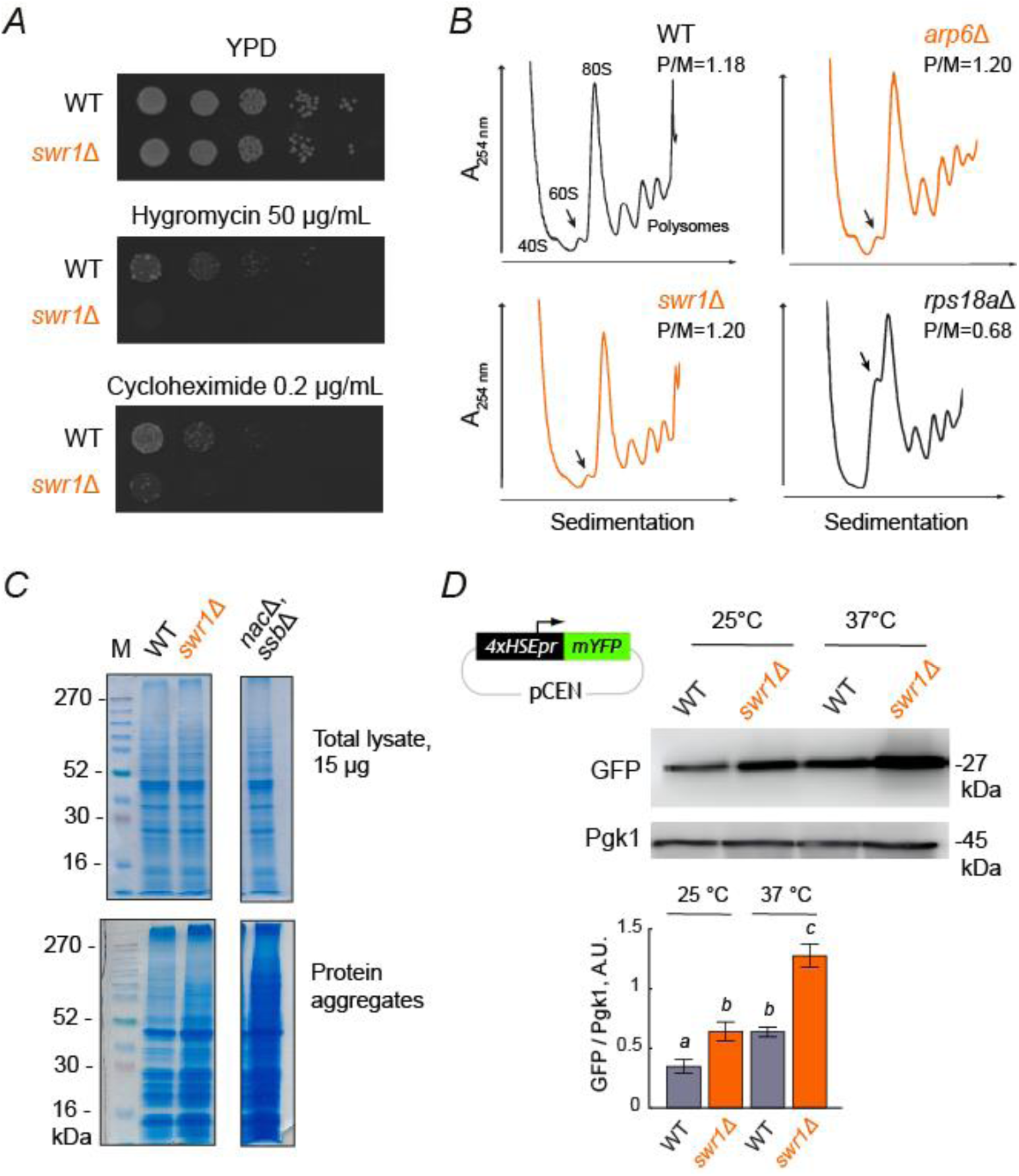
Loss of SWR1C perturbs translation and triggers stress responses. (*A*) WT and *swr1*Δ cells were tested for sensitivity to translation inhibitors. Log-phase cultures were serially diluted (1:10) and spotted onto YPD medium with or without hygromycin B or cycloheximide. Plates were incubated at 30 °C for 48‒72 h before imaging. Images represent three independent experiments with consistent results. **(*B*)** Polysome profiles from WT and gene-deletion strains. Cells were grown to mid-log phase, lysed, and loaded onto linear sucrose gradients; ribosomal species (40S, 60S, 80S, polysomes) were visualized by absorbance at 254 nm. The average polysome-to-monosome ratio (P/M) was calculated from the profile’s AUC. Profiles are representative of three independent biological replicates. **(*C*)** Detection of aggregated proteins in WT and *swr1*Δ mutants. Cells were grown to logarithmic phase in SC medium at 30 °C and lysed; soluble and aggregated fractions were analyzed by SDS-PAGE and visualized by Coomassie staining. Total protein from whole-cell lysates (top) and aggregated fractions (bottom) was loaded. **(*D*)** Hsf1 activity reporter assay. WT and *swr1*Δ strains transformed with a centromeric pRS316 4xHSEpr-mYFP (GFP) plasmid and grown in SD medium at 25 °C following heat shock for 1 h at 37 °C. Protein extracts were analyzed by Western blot using Pgk1 as a loading control and quantified by densitometry analysis. Bar plots (bottom) show mean densitometry ±S.E.M (*n*=3); distinct letters indicate groups that differ significantly (*p*<0.05, one-way ANOVA with Tukey’s post hoc test).

### 2.4 SWR1C impairment drives increased protein aggregation and activates the Hsf1 response

Translation fidelity and efficiency are generally altered in aged cells across yeast, mice, and humans (Gerashchenko et al. 2021; Gonskikh and Polacek 2017; Hansen et al. 2007; Hu et al. 2018; Tuller et al. 2010). In line with our findings, Stein and colleagues showed that aging modifies the kinetics of translation elongation without affecting global pausing translation rates in yeast and worms (Stein et al. 2022). Based on these observations, we hypothesized that the absence of *SWR1* promotes lifespan extension by impairing nascent peptide elongation at the ribosome, leading to protein misfolding and activation of proteostatic stress responses. To test whether *swr1Δ* cells accumulate misfolded proteins, we isolated insoluble protein aggregates from logarithmically growing cultures and analyzed them by SDS-PAGE followed by Coomassie staining. Our results revealed a modest increase in aggregate accumulation in *swr1*Δ cells compared to WT (**Figure 3C**), yet this effect was weaker than that observed in the reference mutant lacking the NAC and Ssb1/2 chaperones, which showed strong aggregate accumulation as expected (Koplin et al. 2010).

Given that Hsf1 is a key transcription factor activated by proteotoxic stress (Kmiecik and Mayer 2022; Tye and Churchman 2021), we next examined whether loss of SWR1 affects Hsf1 activity using a reporter based on a GFP variant driven by a synthetic promoter containing four tandem heat-shock elements (4×HSE), which are specifically responsive to Hsf1. Deletion of *SWR1* led to increased reporter expression both at the permissive temperature (25°C) and, more prominently, under heat-shock conditions (37°C), consistent with elevated proteotoxic stress in these cells (**Figure 3D**). Together, these findings support a model in which loss of SWR1C promotes longevity by inducing mild proteotoxic stress through altered translation dynamics, thereby activating adaptive stress response pathways that enhance survival of aging cells.

### 2.5 Lifespan profiling links non-coding RNAs to aging in *S. cerevisiae*

SWR1C has been reported to regulate non-coding transcription, particularly at repressed loci (Alcid and Tsukiyama 2014), and the histone variant H2A.Z is critical for maintaining the chromatin architecture required for proper tRNA gene expression (Mahapatra et al. 2011). Given our findings that *swr1*Δ mutants display lifespan extension linked to proteostasis without altering global translation rates, we hypothesized that SWR1C may influence longevity through its impact on non-coding RNAs, particularly those involved in translation. Altered tRNA expression could compromise translational fidelity and elongation dynamics, thereby contributing to the mild proteotoxic stress observed in SWR1C-deficient cells.

To test this hypothesis, we first examined whether non-coding RNAs influence aging using CLS assays on a comprehensive deletion library of ncRNA loci (Parker et al. 2017, 2018). This collection was tagged with mCherry (RFP) using the previously described SGA-based method (**Figure S2**), enabling high-resolution competitive CLS screening. The ncRNA deletion library contained 130 tRNA deletions, 62 stable untranslated transcripts (SUTs), 35 cryptic unstable transcripts (CUTs), and 43 small nucleolar RNAs (snoRNAs) (**Dataset S2**). Relative lifespan (*L*_xΔ_) was measured for all mutants in duplicate, yielding highly reproducible results (*r*=0.91; **Figure S3B**).

The frequency distribution of relative lifespan values across all deletion mutants revealed that a remarkable fraction of ncRNA genes modulate CLS in yeast (**Figure 4A**; **Dataset S2**). To score ncRNA-deletion strains with increased or decreased CLS, the distribution of survival coefficients from all deletion strains was compared to a null distribution derived from WT strain replicates. Using a 5% FDR threshold, 27.8% of ncRNA-gene deletions (75/270) produced CLS phenotypes, with a pronounced overall bias toward reduced lifespan (62 short-lived vs 12 long-lived strains), indicating that loss of ncRNA function frequently compromises post-mitotic survival. This overall occurrence of CLS phenotypes among ncRNA-gene deletions was significantly higher than that reported for protein-coding genes (*p*=0.002; Fisher’s exact test). Specifically, applying the same CLS framework and significance threshold to previous large-scale analyses of non-essential coding genes yielded phenotypes in only 20.1% (778/3878) of mutants (Garay et al. 2014), highlighting ncRNA genes as major determinants of chronological longevity.

**Figure 4.**
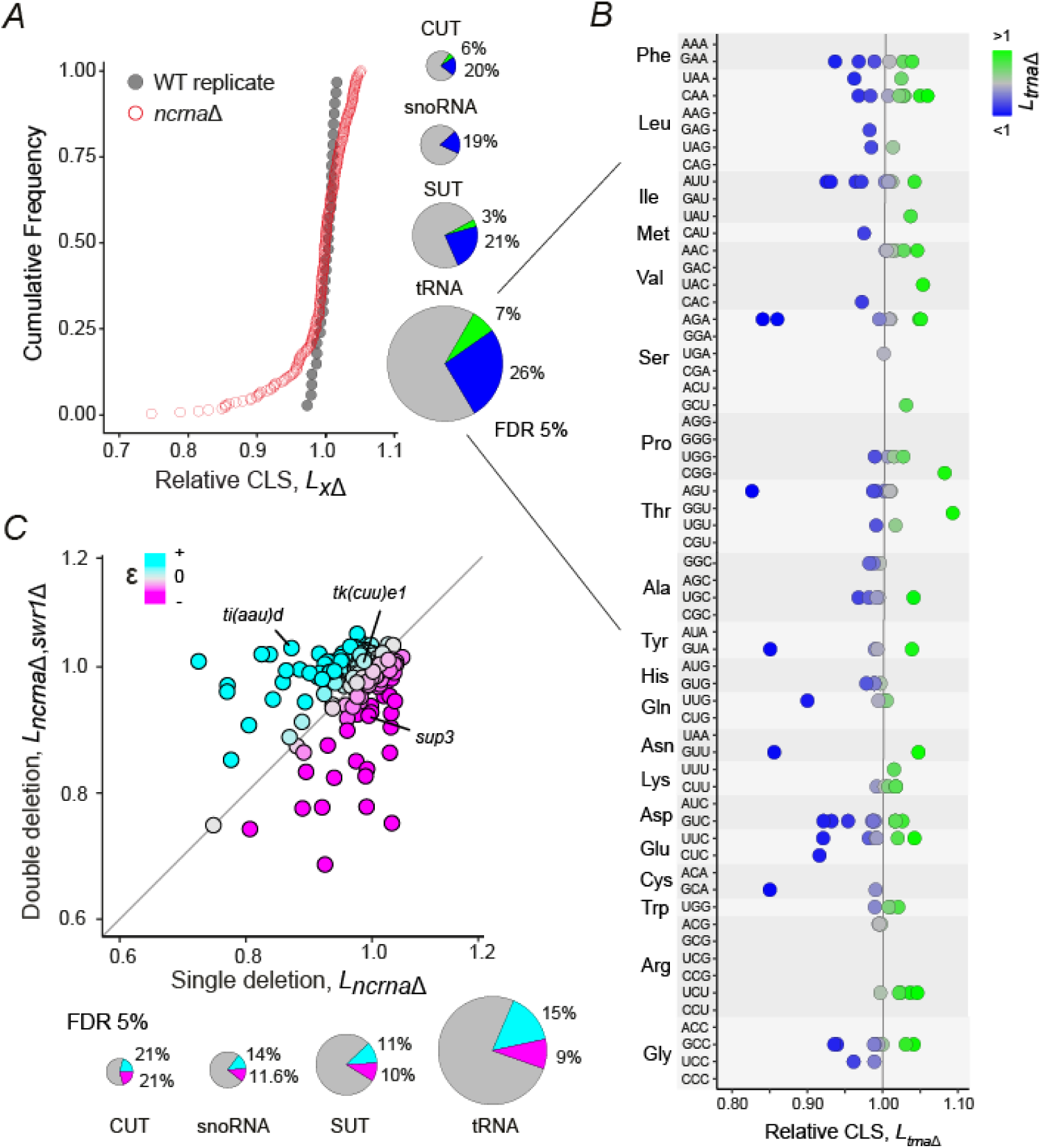
Deletion of tRNAs reveals lifespan phenotypes that are functionally linked to *SWR1*. (*A*) Cumulative frequency distribution of relative CLS (*L_x_*_Δ_) values for ncRNA deletion strains (red open circles, *n*=270) and WT replicates (gray closed circles, *n*=32). *Right:* Pie charts show the fraction of long-lived (green) and short-lived (blue) strains within each ncRNA category. Lifespan phenotypes were classified using a 5% FDR cutoff (Benjamini-Hochberg) based on the reference distribution of WT replicates. **(*B*)** Plot shows the CLS (*L_x_*_Δ_) values of all surveyed tRNA deletions strains (*n*=130), grouped by amino acid isoacceptors and anticodon isodecoders. **(*C*)** Scatter plot showing the CLS *L_x_*_Δ_ values of 259 single-deletion strains compared to their *swr1*Δ double mutants relative to the *swr1*Δ (*L_x_*_Δ*swr1*Δ_). Color scale indicates the epistatic interaction value (ɛ), directly calculated as the difference between the two measurements. *Bottom*: Pie charts summarize the fraction of positive (cyan) and negative (magenta) epistatic interactions within each ncRNA category; interactions were classified with at 5% FDR with 31 WT-reference replicates.

At the 5% FDR threshold used, CLS effects were detected across all ncRNA classes and were generally biased toward reduced lifespan. CUT deletions showed significant phenotypes in 9 of 35 strains, while snoRNA and SUT deletions were associated with CLS changes in 8 of 43 and 15 of 62 strains, respectively (**Figure 4A**, *right*). Importantly, tRNAs, as the largest ncRNA class tested, contributed the greatest absolute number of CLS-altering deletions (52 of 130 tRNAs, 40.0%), with the majority of significant effects also corresponding to reduced lifespan. This finding underscores the exceptional sensitivity of chronological longevity to perturbations in the tRNA pool.

In this context, the ncRNA deletion collection included 47.3% (130/275) of the annotated *S. cerevisiae* tRNA gene repertoire, with multiple isoacceptors and isodecoders for most amino acids. Despite this apparent redundancy, we observed wide variation in lifespan effects among tRNA genes belonging to the same isotype (**Figure 4B**). For example, deletion of the threonine tRNA gene *tT(AGU)C* resulted in a pronounced short-lived phenotype, whereas deletion of five other *tT(AGU)* isodecoder genes had little or no effect on CLS. Conversely, deletion of the threonine isoacceptor *tT(UGU)P* produced the strongest long-lived phenotype observed. Likewise, deletions of *SUP3* or *SUP8*, both encoding *tY(GUA)* tyrosine tRNAs, had no effect on CLS, whereas the *SUP5* and *SUP7* isodecoders were associated with significantly short– and long-lived phenotypes, respectively. Overall, the absolute CLS effects (|*S*|) of tRNA deletions did not correlate with codon usage bias (*r*=‒0.09, *p*=0.33, Pearson). Together, these findings indicate that genetically distinct tRNA genes encoding the same amino acid and even sharing the same anticodon are not fully redundant and contribute differentially to aging, likely due to differences in expression levels or regulatory crosstalk.

### 2.6 Pervasive epistasis between *SWR1* and tRNA genes shapes lifespan outcomes

Given our finding that specific non-coding RNAs strongly influence yeast chronological aging, we asked whether these lifespan effects are functionally linked to SWR1C. To address this, we systematically quantified epistatic interactions between *SWR1* and ncRNA genes by measuring the CLS of the corresponding single and double mutants. For increased resolution, the *x*Δ*swr1*Δ double-mutant array was competed against a *swr1*Δ single mutant, providing a direct readout of epistasis by comparing each deletion’s effect in the WT background and in the *SWR1*-impaired background (**Figure 4C**; **Dataset S3**). Loss of *SWR1* either exacerbated or alleviated the phenotype of the corresponding ncRNA mutant, a pattern that held across all ncRNA categories (**Figure 4C**, *bottom*). Notably, tRNAs accounted for the largest fraction of positive epistatic interactions, in which the deleterious CLS effect of the ncRNA deletion was alleviated by the absence of *SWR1*. This mirrors the trend observed for *SWR1* interactions with genes involved in cytosolic translation, rRNA biogenesis, and proteostasis (**Figure 2B**), further underscoring the close functional relationship between SWR1C and protein homeostasis.

To more precisely characterize the epistatic relationships between *SWR1* and tRNA genes, we selected a subset of 25 tRNA deletions and re-evaluated them in a focused competitive-aging screen with increased replication. This subset primarily included tRNA deletions with a detectable CLS phenotype, along with a few isoacceptors and isodecoders with neutral effects. This design enabled accurate scoring of negative, positive, and neutral epistasis based on deviations of double-deletion CLS from the multiplicative expectation derived from the corresponding single mutants (**Figure 5A**). For example, deletion of the isoleucine tRNA gene *tI(AAU)D* caused a strong reduction in lifespan relative to WT, which was fully suppressed in the *swr1*Δ*tI(aau)d*Δ double mutant. Conversely, deletion of the tyrosine isodecoder *tY(GUA)O* (*SUP3*) had no effect on CLS alone but led to a markedly shorter lifespan when combined with *swr1*Δ. These experiments confirmed widespread genetic interactions between tRNA genes and *SWR1*, with many double mutants showing reduced lifespan relative to single deletions, indicative of negative epistasis (**Figure 5B**). Specifically, 44% (11/25) of the tRNAs surveyed showed a significant aggravating interaction with *swr1*Δ, whereas 12% (3/25) displayed positive epistasis (**Figure S4**). This pattern reflects the particular subset examined, confirming that SWR1C-dependent longevity is sensitive to perturbations in specific tRNA genes. Together with the broader analysis, these findings support a role for tRNAs as key modulators of the longevity phenotype associated with *SWR1* loss.

**Figure 5.**
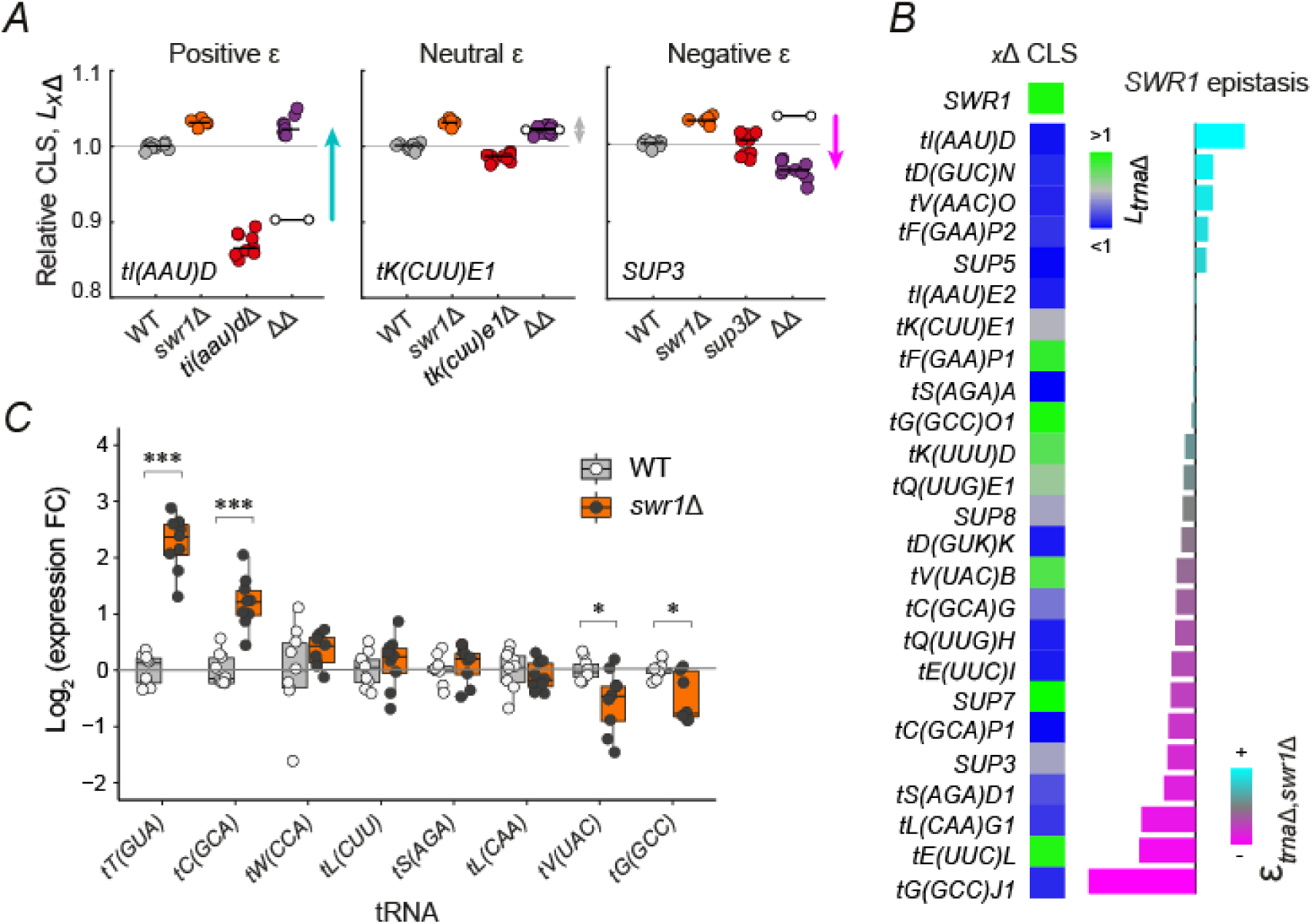
Deletion of *SWR1* modifies the CLS mutant phenotype and expression of tRNA genes. (*A*) Examples of genetic interactions between *SWR1* and tRNA mutants. Plots show the measured relative CLS (*L*_xΔ_) of single and double knockouts. Solid black lines indicate the mean of 9 replicates for each tRNA mutant and 6-10 replicates for WT and *swr1*Δ in each comparison. Circle-endpoint lines (⧟) show the expected multiplicative lifespan (*L_x_*_Δ_)·(*L_swr1_*_Δ_) for the *L_x_*_Δ*swr1*Δ_ double mutant. Significant deviations from the expected are positive (cyan) or negative epistasis (magenta) (*p*<0.05; *t*-test). **(*B*)** The CLS of tRNA mutants is usually altered in the *swr1*Δ background. The heatmap shows CLS values for selected tRNA mutants, with color scale indicating long-lived (green), neutral (gray), and short-lived (blue) phenotypes. Adjacent horizontal bar plot shows the distribution of average epistatic interactions across nine replicates per gene pair (**Figure S4**). Color scale indicates positive (cyan), neutral (gray), and negative (magenta) epistasis. **(*C*)** Deletion of *SWR1* alters tRNA abundance. Expression fold-changes (FC) of individual tRNAs were measured by RT-qPCR and calculated using the 2^−ΔΔCT^ method, based on cycle threshold (CT) values relative to the WT. PCR primers for accurate tRNA monitoring were from (Torrent et al. 2018) with *ACT1* used as an internal normalization control. Bars represent the mean ± S.E.M. of three biological replicates, each performed in triplicate. Statistical significance is indicated (\**p*<0.05, \*\*\**p*< 0.001; *t*-test).

To directly test whether SWR1C affects CLS through changes in tRNA pool composition, we focused on tRNAs whose deletions genetically interacted with *swr1*Δ and measured the expression of eight such tRNAs in the *SWR1*-deficient background. We employed a modified RT-qPCR approach, allowing us to monitor processed tRNA species with enhanced sensitivity (Torrent et al. 2018). Because many tRNAs are encoded by multiple genomic copies with nearly identical sequences, these RT-qPCR measurements reflect the combined abundance of all processed transcripts amplified by each primer pair. In *swr1*Δ strains, we observed strong increased expression of two tRNAs, specifically *tY(GUA)* and *tC(GCA)*, along decreased expression of *tV(UAC)* and *tG(GCC)* (**Figure 5C**). These changes suggest a substantial shift in the relative abundance of tRNAs available for translation in SWR1C-impaired cells. Our results indicate that *SWR1* loss alters the expression of specific tRNAs, modulating the lifespan phenotypes associated with individual tRNA deletions. Given established links between tRNA availability, translational efficiency, and proteostasis (Gingold and Pilpel 2011; Percudani et al. 1997; Tuller et al. 2010), these findings support a model in which altered tRNA pool composition contributes to the longevity phenotype of *swr1*Δ cells.

### 2.7 Longevity in *swr1Δ* cells is associated with tRNA-mediated ER proteostasis surveillance

To determine how altered tRNA pools in *swr1*Δ cells affect proteostasis pathways, we examined markers of ER stress and protein-folding surveillance. We focused on the tY(GUA) genes, which encode tyrosine isodecoders, as they showed strong genetic interactions with SWR1 and increased overall expression in the *swr1*Δ background. To assess proteotoxic stress sensitivity, we performed serial dilution spot assays using tunicamycin, which induces protein unfolding by inhibiting ER glycosylation (**Figure 6A**). Loss of *SWR1* resulted in a subtle increase in tunicamycin tolerance, consistent with a minor shift in ER stress sensitivity. A clear increase in ER-stress tolerance was observed in the *sup3Δ* and, more prominently, *sup7*Δ tRNA mutants. Notably, this enhanced tolerance was abolished—and even reversed—in the *sup3*Δ*swr1*Δ and *sup7*Δ*swr1*Δ double mutants, which exhibited increased sensitivity to tunicamycin-induced ER stress. By contrast, the *sup5*Δ single mutant displayed strong tunicamycin sensitivity, which was fully suppressed in the Δ*sup5*Δ*swr1* double mutant. No changes in ER-stress sensitivity were observed in strains lacking *SUP8*, either alone or in combination with *swr1*Δ. Together, these patterns indicate that tyrosine-decoding tRNAs modulate the ER-stress phenotype associated with *SWR1* loss.

**Figure 6.**
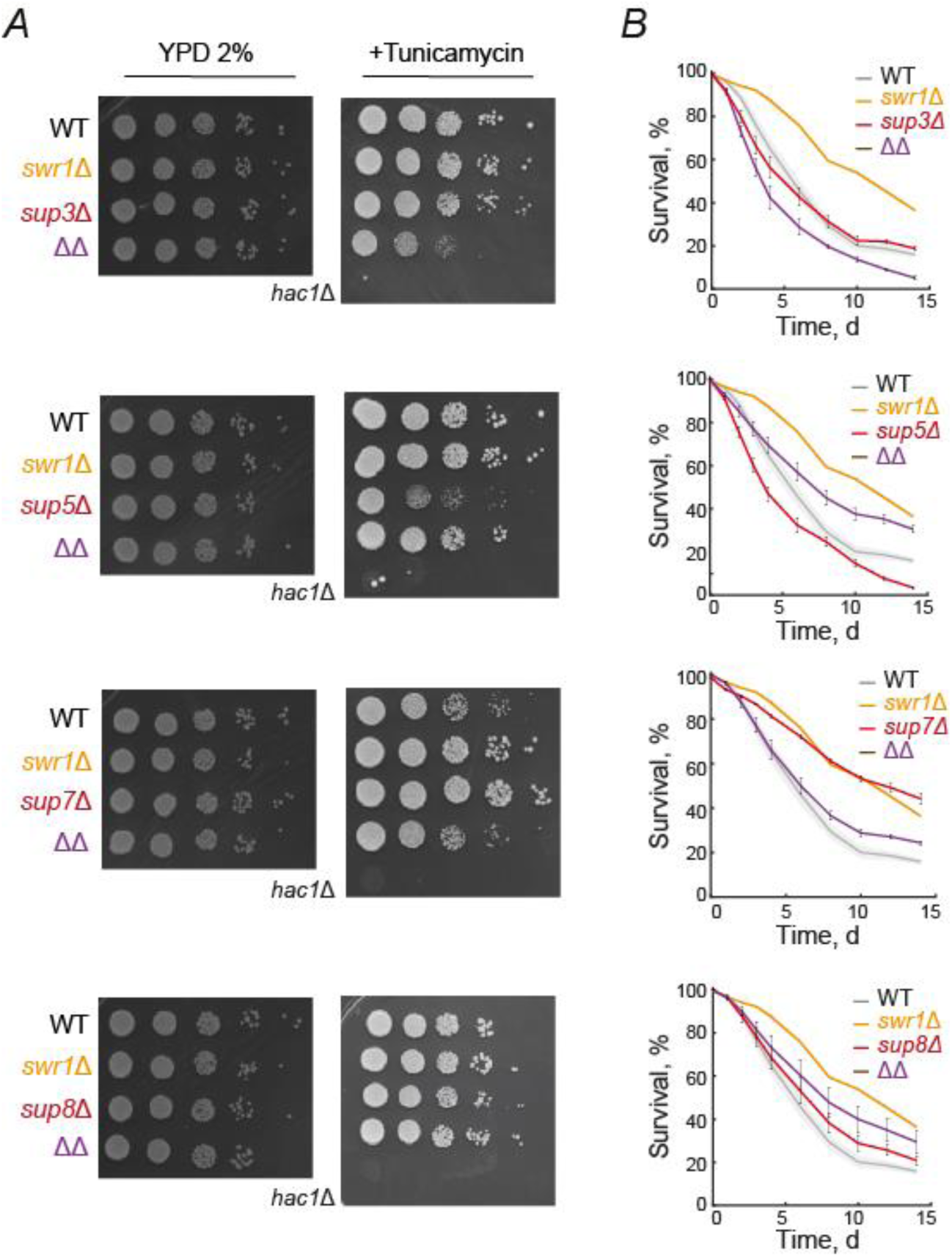
Functional interactions between *SWR1* and tyrosine tRNAs modulate ER proteostasis surveillance and impact cellular lifespan. (*A*) WT, single– and double-gene deletion strains were cultured in SC medium to an OD_600_ of 0.7. Serial 1:10 dilutions were spotted onto YPD plates, either alone or supplemented with 0.75 μg·mL⁻¹ tunicamycin. Plates were incubated at 30 °C for 48 hours before imaging. All experiments were performed in three independent replicates with consistent results. (*B*) CLS was assessed using a live/dead staining assay. Survival curves show WT (gray), *swr1Δ* (orange), single *tRNA-Tyr* deletions (red), and double mutants (purple) over time. Shaded areas and error bars represent the mean ± S.E.M from four independent replicates (*n*=4).

To determine whether these ER-linked *SWR1*–tRNA interactions directly translate into aging phenotypes, we evaluated their effects on CLS (**Figure 6B**). This analysis revealed that the reduced lifespan of the *sup3*Δ*swr1*Δ and *sup7swr1*Δ double mutants relative to their corresponding single mutants mirrors their increased sensitivity to ER stress. Likewise, the suppression of strong tunicamycin sensitivity in the *sup5*Δ mutant by *swr1*Δ was recapitulated at the level of lifespan, with the double mutant exhibiting improved CLS. One exception was *SUP8*, where the double mutant displayed a shorter lifespan than expected based on the single mutants, despite showing no change in ER-stress sensitivity, suggesting that additional factors beyond proteostasis contribute to the observed aging phenotypes. Considering the higher identity of the four genes surveyed encode tyrosine *tY(GUA) tRNAs*, these findings demonstrate that SWR1C modulates the functional output of specific tRNAs, with consequences for protein homeostasis that ultimately shape lifespan outcomes.

## 3 Discussion

Aging is shaped by both nuclear and cytoplasmic pathways, yet the mechanisms that connect chromatin remodeling to proteostasis remain obscure. SWR1C, a conserved chromatin-remodeling complex best known for mediating H2A.Z exchange, transcriptional regulation, and genome stability (Van Attikum et al. 2007; Kobor et al. 2004; Wu et al. 2005), has been reported to extend chronological lifespan when disrupted in both *S. cerevisiae* (Garay et al. 2014) and *S. pombe* (Romila et al. 2021). However, how SWR1C influences longevity has remained unresolved. Here we show that loss of SWR1C components—including multiple subunits, H2A.Z, and *SWR1* across divergent *S. cerevisiae* backgrounds—consistently extends lifespan, revealing SWR1C as a conserved pro-aging factor in ascomycete yeasts. Importantly, our genetic and cellular analyses uncover an unexpected connection between SWR1C impairment, tRNA homeostasis, and proteostasis capacity, establishing a mechanistic link between chromatin remodeling, tRNA biology, and aging.

The widespread conservation of H2A.Z and *SWR1*-like remodelers suggests that related regulatory circuits linking chromatin dynamics to tRNA biology and proteostasis may operate in other eukaryotes. Global reductions in tRNA synthesis, such as those produced by Pol III inhibition, enhance resistance to proteostasis stress and extend lifespan across species (Hansen et al. 2007; Malik et al. 2024), underscoring tRNA homeostasis as a conserved determinant of aging. Our findings refine this framework by showing that SWR1C does not act as a broad repressor of Pol III activity but instead modulates tRNA abundance at the level of specific gene copies. This selective remodeling of the tRNA pool aligns with prior evidence that tRNA genes within the same isotype can have distinct physiological roles (Sagi et al. 2016) and extends this principle to lifespan regulation. Moreover, we identified tyrosine tRNA isodecoders as key modifiers of the *swr1*Δ aging phenotype, supporting a model in which defined tRNA changes reshape translational homeostasis and proteostasis with measurable impacts on longevity.

Unlike interventions that extend lifespan through broad translational repression (Hansen et al. 2007; Howard et al. 2016), *swr1*Δ cells retain largely unaltered polysome profiles and show no evidence of global defects in translation initiation or elongation. Instead, they display increased sensitivity to translation inhibitors, mild accumulation of protein aggregates, and activation of the Hsf1 regulon. This combination of phenotypes suggests a reorganization of translational fidelity and proteostasis capacity rather than reduced translation output. Notably, changes in tRNA pool composition could generate codon-specific ribosome pauses that impact co-translational folding, a process that is exacerbated with age in budding yeast and nematodes (Stein et al. 2022). Whether altered tRNA availability in *swr1*Δ cells directly modulates ribosome pausing or mRNA-specific translation elongation dynamics remains to be addressed. The robust heat-shock response further suggests that SWR1C loss triggers a compensatory proteostasis program that supports survival under stationary-phase stress. Together, these observations position SWR1C as a regulator of the balance between translational accuracy, folding stress, and protein turnover rather than a mediator of classical metabolic or stress-resistance pathways.

The association between SWR1C, H2A.Z exchange, and tRNA regulation adds a new layer to our understanding of chromatin-ncRNA interplay. Lifespan extension correlated strictly with deletions of SWR1C subunits required for H2A.Z deposition, indicating that the mechanism depends on impaired histone variant exchange rather than indirect stress responses. This aligns with prior evidence that SWR1C-mediated H2A.Z turnover modulates antisense and noncoding transcription (Alcid and Tsukiyama 2014). Although our study did not include genome-wide chromatin profiling, earlier analyses detected H2A.Z enrichment and altered nucleosome organization near Pol III loci (Albert et al. 2007; Yen et al. 2013), suggesting that SWR1C disruption may alter chromatin accessibility at specific tRNA genes. Consistent with this, targeted quantification of selected tRNAs revealed changes in abundance that align with lifespan-genetic interactions, supporting a model in which chromatin state influences tRNA gene regulation and, in turn, translational homeostasis.

This study uncovers an unanticipated role for the SWR1C chromatin-remodeling complex in lifespan regulation through selective modulation of tRNA abundance and proteostasis. The observation that individual tRNA genes exert distinct and epistatic effects on longevity reveals a previously unappreciated layer of specificity in chromatin-mediated control of cellular homeostasis and positions SWR1C as a key modulator of translational integrity. Although the chromatin mechanisms underlying these selective tRNA changes remain to be resolved, future analyses of nucleosome organization and Pol III-associated regions (Albert et al. 2007; Yen et al. 2013) will help clarify how SWR1C remodeling shapes tRNA gene regulation. More broadly, our findings provide a mechanistic framework for how chromatin remodelers influence aging through RNA-based regulation of translation and proteostasis, with potential relevance to age-associated proteostasis disorders.

## 4 Materials and Methods

### 4.1 Strains and media

Yeast single mutants were generated using the standard LiAc-PEG transformation method (Gietz 2014). Gene-deletion strains *swr1*Δ, *arp6*Δ, *swc5*Δ, *swc7*Δ, *swc6*Δ, *swc2*Δ, *bdf2*Δ, *yaf9*Δ, *pho23*Δ, *eaf3*Δ, *eaf7*Δ, *eaf6*Δ, *htz*1Δ, and the neutral *his3*Δ and *ho*Δ insertions were constructed de novo in the YEG01-RFP background (Garay et al. 2014) by PCR-based replacement with *NatMX6* from pAG25 (Euroscarf). All knockouts were PCR-verified for loss of the target ORF and correct marker integration. The resulting genotype was *Mat*α *PDC1-mCherry-CaURA3MX4, can1*Δ::*STE2pr-SpHIS5*, *lyp1*Δ, *his3*Δ*1*, *ura3*Δ*0*, *LEU2*, *x*Δ::*NatMX6*, with *his3*Δ or *ho*Δ used as the WT reference strains. The *swr1*Δ and *ho*Δ alleles in strains YJM981, LY374, Y12, and YJM978 (Liti et al. 2009) were similarly generated by homologous recombination.

Growth media included: (1) aging medium (SC with 2% dextrose, 0.2% amino acid mix; Sigma Y1501, supplemented with uracil; Sigma U0750, unbuffered); (2) low-fluorescence medium (YNB-lf; DeLuna et al., 2008); and (3) YPD or YPAD (2% dextrose). Detailed media compositions are provided in **Table S2**.

### 4.2 RFP-tagged yeast deletion libraries

A set of 314 viable single-gene deletion strains previously linked to chronological aging pathways (Cruz-Bonilla et al. 2025) was selected from the yeast deletion collection, all in the BY4741 background (*MAT*a *xxxΔ::kanMX4 his3Δ1 ura3Δ0 leu2Δ0 met15Δ0*) (**Dataset S1**). These strains were crossed to the *Mat*α YEG01-RFP *swr1*Δ::*NatMX6* SGA-starter strain methodology (Tong and Boone 2006) (**Figure S2**). Crosses were performed on 384-colony arrays transferred manually with a VP384F headpin tool (V&P Scientific). Standard SGA steps—mating, diploid selection, sporulation, and three rounds of haploid selection—resulted in isogenic double mutants with the genotype *Mat*a *xxxΔ::kanMX4 swr1Δ::NatMX6 PDC1*-*mcherry*-*CaURA3MX4 can1Δ:STE2pr*-*SpHIS5 lyp1Δ his3Δ1 ura3Δ0 LEU MET LYS*. Antibiotic concentrations used for selection were 200 µg·mL⁻¹ G418 (Invitrogen), 100 µg·mL⁻¹ clonNAT (Werner BioAgents), 100 µg·mL⁻¹ canavanine, and 100 µg·mL⁻¹ thialysine (Sigma-Aldrich).

For CLS screening of non-coding RNA genes, the collection described by (Parker et al., 2018) was used as the basis for generating an RFP-tagged ncRNA knockout set. Strains were tagged using the SGA procedure described above and in **Figure S2**. The resulting genotype was *Mat a xncrnaΔ::kanMX4 PDC1*-*mcherry*-*CaURA3MX4 can1*Δ*:STE2pr*-*SpHIS5 lyp1*Δ *his3*Δ*1 ura3*Δ*0 LEU MET LYS* (**Dataset S2**). The same approach was used to generate the RFP-tagged *swr1*Δ ncRNA double-deletion collection (**Dataset S3**).

### 4.3 Automated competitive-aging screens and data analysis

Overnight cultures of RFP-tagged gene or ncRNA deletion strains into 96-well plates (150 µL aging medium) and reference strains were grown separately and then mixed at a 2:1 ratio. For CLS screening, mixed cultures (∼1.5 µL) were transferred with a 96-pin tool (V&P VP407) into 96 semi-deepwell plates containing 700 µL aging medium (stationary phase cultures) and incubated at 30 °C without shaking. Each plate included 8–10 reference competitions (WT-RFP/WT-CFP or *swr1*Δ-RFP/*swr1*Δ-CFP). Detector gains were optimized (CFP 140–160; RFP 240–260). At 72 h (stationary phase, time zero) and subsequently every 24–48 h for up to 10 days, Aging Cultures were resuspended and 10 µL were transferred robotically into 140 µL YNB-lf to generate outgrowth cultures. RFP (Ex 587/5, Em 610/5), CFP (Ex 433/5, Em 475/5), and OD600 were measured every 90 min for ∼14 h using a Tecan Freedom EVO200–M1000 system, with shaking before each reading.

The natural log of the RFP/CFP ratio across outgrowth time points was fitted to a multiple linear model to derive the survival coefficient (*S*) and relative CLS (*L* = 1 + *S*). The CLS-profiling workflow is shown in **Figure 1C**; model and data fitting details are provided in (Avelar-Rivas et al. 2020).

### 4.4 Scoring CLS phenotypes and lifespan-epistasis interactions

To identify short– and long-lived mutants in single– and double-deletion screens, *Z*-scores were computed for each experimental *S* value relative to an empirical null distribution defined by WT reference strain replicates (WT-RFP/WT-CFP competition). The null distribution was characterized by its mean (μ) and standard deviation (σ), estimated from all reference measurements. Each experimental value was then standardized as:

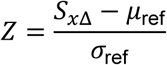

Two-sided *p*-values were calculated from the standard normal distribution. To account for multiple testing, *p*-values were adjusted using the Benjamini–Hochberg false discovery rate (FDR) procedure. Mutants with an FDR-adjusted *p*-value ≤ 0.05 were considered to exhibit significant short-or long-lived phenotypes (5% FDR cutoff).

Epistasis (ε) was defined as the difference between double– and single-mutant *S* values. Importantly, single-mutant *S* values were obtained from competitive aging assays using *x*Δ-RFP/WT-CFP strains, whereas double-mutant *S* values were obtained from the *x*Δ*swr1*Δ-RFP/*swr1*Δ-CFP competition. Epistasis was calculated directly as:

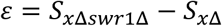

Epistasis values were subsequently used for functional enrichment analyses.

### 4.5 Functional GO-enrichment analysis

The median epistasis (ɛ) of genes associated with each Gene Ontology (GO) term was compared to the median epistasis of the entire dataset. Genes were grouped according to their annotated GO terms, with only groups containing at least five and no more than 50 genes being considered. Annotation files were obtained from the Yeast Genome Database (2024). Go terms with significantly different ɛ median values (*p*<0.05, Wilcoxon rank sum test) are shown (**Figure 2B**; **Table S1**).

### 4.6 CLS assays by live/dead staining

Monocultures were grown to stationary phase in semi-deepwell plates. At day 2 and subsequently up to day 14, 6 µL of culture were transferred to 50 µL BD FACSFlow (BD 342003) containing LIVE/DEAD™ FungaLight™ (ThermoFisher L34952) in 96-well plates. Samples were shaken for 1 min at 1,000 rpm (Heidolph Titramax 1000) and incubated 20 min at room temperature in the dark. PI and SYTO9 fluorescence were measured on an LSRFortessa™-HTS (BD), acquiring 10,000 events per sample. Cells with high SYTO9 and low PI fluorescence were scored as alive, whereas PI-positive cells were classified as dead. The percentage of live cells at each time point was used to generate survival curves. Survivorship at each time point was calculated as the mean percentage of live cells across replicates. For statistical comparison, the AUC from each individual replicate was used, and differences relative to WT were assessed by a two-tailed *t*-test.

### 4.7 Serial dilution spot assays

Selected strains were grown overnight in aging medium, diluted to OD_600_=0.3, and incubated at 30 °C with shaking until OD600 reached 0.7–1.0. Cultures were normalized to OD_600_=1.0, pelleted (10,000 rpm), and resuspended in 1 mL water. Fivefold serial dilutions were prepared and 3 µL of each were spotted onto YPD plates supplemented with 50 µg·mL⁻¹ hygromycin, 0.3 µg·mL⁻¹ cycloheximide, or 0.75 µg·mL⁻¹ tunicamycin. Plates were incubated at 30 °C for 48–72 h and photographed. All assays were performed with at least three independent replicates.

### 4.8 Polysome profiling

WT and mutant strains were grown in 100 mL YPD at 30 °C to OD_600_ 0.5-0.7. Cycloheximide was added to 100 µg·mL⁻¹, cultures were rapidly chilled for 15 min, and cells were harvested at 3,500 g for 10 min (Beckman JA-14). Pellets were washed twice with LHB buffer (10 mM Tris-HCl pH 7.4, 100 mM NaCl, 30 mM MgCl_2_, 500 µg·mL⁻¹ cycloheximide, 200 µg·mL⁻¹ heparin), resuspended in 0.5 mL LHB with protease inhibitors, and lysed with 500-µm glass beads in a FastPrep-24™ (five 30-s cycles with 3-min cooling). Lysates were supplemented with 1 mL cold LHB and clarified twice at 10,000 g for 10 min at 4 °C. Ninety A_260_A units were loaded onto 10-35% sucrose gradients in TMN buffer (50 mM Tris/acetate pH 7.0, 50 mM NH_4_Cl, 12 mM MgCl_2_) and centrifuged at 25,000 rpm for 5.5 h at 4 °C (SW28 rotor). Gradients were fractionated with continuous A₂₅₄ monitoring. From the resulting polysome profiles, the areas under the 80S monosome peak and the polysomal peaks were calculated to derive the polysome-to-monosome (P/M) ratio.

### 4.9 RNA isolation, cDNA synthesis, and qRT-PCR for tRNA expression analysis

Total RNA was isolated following the TRIzol phenol-chloroform protocol described in (Torrent et al. 2018). Five milliliters of logarithmic-phase culture (OD_600_=0.5) were harvested at 3,000 g for 3 min, washed with nuclease-free water, and pelleted again at 3,000 g. Cells were resuspended in 150 µL lysis buffer (0.1 M lithium acetate, 0.5% SDS) and incubated at 70 °C for 5 min. TRIzol LS® (450 µL) was added and mixed for 15 s, followed by 150 µL chloroform and another 15-s mix. After 5 min at room temperature, samples were centrifuged at 12,000 g for 30 min at 4 °C. The aqueous phase was transferred to 450 µL isopropanol and RNA was precipitated overnight at –20 °C. RNA was collected at 10,000 g for 20 min at 4 °C, washed three times with 75% ethanol, air-dried for ≥1 h, and dissolved in 20 µL nuclease-free water (Qiagen). RNA quantity was determined with a NanoDrop ND-1000, ensuring A260/230 and A260/280 ratios > 2. RNA integrity was assessed on 1% agarose gels using 500 ng RNA.

RNA samples were treated with DNase I (New England Biolabs) according to the manufacturer’s instructions. cDNA was synthesized from 1 µg total RNA using the RevertAid First Strand cDNA Synthesis Kit (Thermo Fisher Scientific) following (Torrent et al. 2018). Expression of selected tRNAs was quantified by qRT-PCR on a StepOnePlus™ Real-Time PCR System (Applied Biosystems). Each reaction contained 10 ng cDNA, SYBR Master Mix (Jena Bioscience), and tRNA-specific primers previously reported by (Torrent et al. 2018) (**Table S3**). Assays were performed in three biological replicates, each in technical triplicate. *ACT1* served as the reference gene. Relative tRNA expression was calculated using the 2^⁻ΔΔCT^ method (Livak and Schmittgen 2001) and reported as mean log^2^ fold-change ±S.E.M. Statistical significance was assessed using a two-tailed Student’s *t*-test (*p*<0.05).

### 4.10 SDS-PAGE and Western Blot Analysis

WT and *swr1*Δ strains carrying the centromeric plasmid pRS316-4xHSE::mYFP::URA2 were grown in SD-Ura to mid-log phase. Cells corresponding to 4 OD_600_ units were harvested and lysed in 50 mM Tris, 0.3 N NaOH, 176 mM β-mercaptoethanol, and 3.5 mM PMSF for 10 min on ice. Proteins were precipitated with 12% (w/v) trichloroacetic acid, washed with cold acetone, and resuspended in 10% SDS and 1X Laemmli buffer. Ten microliters of each extract were separated on 16% SDS–polyacrylamide gels, transferred to nitrocellulose membranes, and probed with anti-Pgk1 (Santa Cruz SC130335) and anti-GFP (Roche 11814460001) antibodies.

### 4.11 Protein aggregation assay

Aggregated proteins were isolated following by (Koplin et al., 2010) with minor modifications. Fifty OD_600_ units of logarithmically growing cells (SC medium) were harvested and resuspended in lysis buffer (20 mM Na-phosphate pH 6.8, 10 mM DTT, 1 mM EDTA, 0.1% Tween, 1 mM PMSF, cOmplete™ Mini protease inhibitors, 3 mg·mL⁻¹ zymolyase, nuclease inhibitors). After 20 min at room temperature, samples were chilled and lysed by tip sonication (Sonics Vibra Cell; five 1-min cycles at 60% amplitude). Lysates were clarified at 200 g for 20 min at 4 °C, and supernatants were normalized to equal protein concentration (Lowry assay). Aggregated proteins (1 mg total protein) were pelleted at 16,000 g for 20 min at 4 °C, washed twice with 2% NP-40 buffer (20 mM Na-phosphate pH 6.8, 1 mM PMSF, protease inhibitors), and centrifuged again at 16,000 g for 20 min at 4 °C. Pellets were washed once in NP-40–free buffer, boiled in 1X Laemmli buffer, resolved on 14% SDS–PAGE gels, and visualized by Coomassie staining.

## Author Contributions

Conceptualization: E.M.-M., A.D. Performed experiments: E.M.-M., J.M.-P., J.U.-C., I.S.-M., L.N.B.-C. Performed data analysis: E.M.-M., A.D. Contributed materials/analysis tools: L.N.B.-C., N.S.P, D.D., K.O., A.D. Supervised experiments and data analysis: N.S.P., D.D., K.O., S.F., A.D. Acquired funding: S.F., A.D. Manuscript writing: E.M.-M., A.D., with input from L.N.B.-C., N.S.P, D.D., K.O., S.F. All authors read and critically revised the final manuscript version.

## Acknowledgements

We are grateful to Diana Ascencio and David Valle-García for critical reading of the manuscript, Cristina Aranda, Arnulfo Bautista, and Ariann Mendoza-Martínez for skillful technical assistance, J. Abraham Avelar-Rivas for valuable help with data analysis, and Erika Garay for useful discussions and contributions to early experimental design.

## Funding

This work was funded by the Secretaría de Ciencia, Humanidades, Tecnología e Innovación de México (Secihti), Grants A1-S-31413 and CF-2023-I-1545. EM-M. was funded by postdoctoral fellowships from the Programa para el Desarrollo Profesional Docente (PRODEP-2019), Secihti (383325), and the Dirección General de Asuntos del Personal Académico (DGAPA-UNAM) (CJIC/CTIC/5642/2022). A.D. was funded by a sabbatical fellowship from Secihti (26244).

## Conflicts of Interest

The authors declare that they have no conflict of interest.

## Data Availability Statement

The data that support the findings of this study are included in this article or available from the corresponding author on reasonable request.

## Contact Information

Ericka Moreno-Mendez

Jimena Meneses-Plascencia

Judith Ulloa-Calzonzin

Ivón Salazar-Martínez

Laura N. Balarezo-Cisneros

Nuria Sánchez-Puig

Daniela Delneri

Katarzyna Oktaba

Soledad Funes

Alexander DeLuna

## Abbreviations

CAN: L-Canavanine
CFP: Cerulean fluorescent protein
CLS: Chronological lifespan
CT: Cycle threshold
CUT: Cryptic unstable transcript
DTT: Dithiothreitol
ɛ: Epistasis
ER: Endoplasmic reticulum
FDR: False discovery rate
GO: Gene ontology
H2A.Z: Histone variant H2A.Z
KanMX: Geneticin resistance cassette
*L*: Relative chronological lifespan
LHB: Label hydrating Buffer
LiAc: Lithium acetate
NatMX: ClonNat resistance cassette
ncRNAs: Non-coding RNAs
OD: Optical density
PEG: Polyethylene glycol
PI: Propidium iodide
RT-qPCR: Reverse transcription quantitative PCR
RFP: Red fluorescent protein (mCherry)
*S*: Survival coefficient
SC: Synthetic complete medium
SD: Synthetic dextrose medium
SDS: Sodium dodecyl sulfate
S.E.M.: Standard error of the mean
SGA: Synthetic genetic array
snoRNA: Small nucleolar RNA
SS: Singel strand DNA carrier from salmon sperm
SUT: Stable untranslated transcript
SWR1C: SWR1 complex
SYTO9: Green-fluorescent nucleic acid stain
THI: Thialysine
TMN: Tris-MgCl2-NaCl buffer
tRNA: Transfer RNA
TSS: Transcription start site
UPR: Unfolding protein response
Ura: Uracil
WT: Wild type strain
YFP: Green fluorescent protein, yellow variant
YNB-lf: Low fluorescence yeast nitrogen base medium
YPAD: Yeast-Peptone-Adenine-Dextrose medium
YPD: Yeast-Peptone-Dextrose medium

## Notes

### Competing Interest Statement

The authors have declared no competing interest.

## References

1. Albert, Istvan, Travis N. Mavrich, Lynn P. Tomsho, Ji Qi, Sara J. Zanton, Stephan C. Schuster, and B. Franklin Pugh. 2007. ‘Translational and Rotational Settings of H2A.Z Nucleosomes across the Saccharomyces Cerevisiae Genome’. Nature 446(7135):572–76. doi:10.1038/NATURE05632.

2. Alcid, Eric A., and Toshio Tsukiyama. 2014a. ‘ATP-Dependent Chromatin Remodeling Shapes the Long Noncoding RNA Landscape’. Genes & Development 28(21):2348–60. doi:10.1101/GAD.250902.114.

3. Alcid, Eric A., and Toshio Tsukiyama. 2014b. ‘ATP-Dependent Chromatin Remodeling Shapes the Long Noncoding RNA Landscape’. Genes & Development 28(21):2348–60. doi:10.1101/GAD.250902.114.

4. Altaf, Mohammed, Andréanne Auger, Julie Monnet-Saksouk, Joëlle Brodeur, Sandra Piquet, Myriam Cramet, Nathalie Bouchard, Nicolas Lacoste, Rhea T. Utley, Luc Gaudreau, and Jacques Côté. 2010. ‘NuA4-Dependent Acetylation of Nucleosomal Histones H4 and H2A Directly Stimulates Incorporation of H2A.Z by the SWR1 Complex’. The Journal of Biological Chemistry 285(21):15966–77. doi:10.1074/JBC.M110.117069.

5. Van Attikum, Haico, Olivier Fritsch, and Susan M. Gasser. 2007. ‘Distinct Roles for SWR1 and INO80 Chromatin Remodeling Complexes at Chromosomal Double-Strand Breaks’. The EMBO Journal 26(18):4113–25. doi:10.1038/SJ.EMBOJ.7601835.

6. Avelar-Rivas, J. Abraham, Michelle Munguía-Figueroa, Alejandro Juárez-Reyes, Erika Garay, Sergio E. Campos, Noam Shoresh, and Alexander DeLuna. 2020. ‘An Optimized Competitive-Aging Method Reveals Gene-Drug Interactions Underlying the Chronological Lifespan of Saccharomyces Cerevisiae’. Frontiers in Genetics 11. doi:10.3389/FGENE.2020.00468.

7. Balarezo-Cisneros, Laura Natalia, Steven Parker, Marcin G. Fraczek, Soukaina Timouma, Ping Wang, Raymond T. O’Keefe, Catherine B. Millar, and Daniela Delneri. 2021. ‘Functional and Transcriptional Profiling of Non-Coding RNAs in Yeast Reveal Context-Dependent Phenotypes and in Trans Effects on the Protein Regulatory Network’. PLoS Genetics 17(1). doi:10.1371/JOURNAL.PGEN.1008761.

8. Bloom-Ackermann, Zohar, Sivan Navon, Hila Gingold, Ruth Towers, Yitzhak Pilpel, and Orna Dahan. 2014. ‘A Comprehensive TRNA Deletion Library Unravels the Genetic Architecture of the TRNA Pool’. PLoS Genetics 10(1):e1004084. doi:10.1371/JOURNAL.PGEN.1004084.

9. Borovinskaya, Maria A., Shinichiro Shoji, Kurt Fredrick, and Jamie H. D. Cate. 2008. ‘Structural Basis for Hygromycin B Inhibition of Protein Biosynthesis’. RNA (New York, N.Y.) 14(8):1590–99. doi:10.1261/RNA.1076908.

10. Campos, Sergio E., J. Abraham Avelar-Rivas, Erika Garay, Alejandro Juárez-Reyes, and Alexander DeLuna. 2018. ‘Genomewide Mechanisms of Chronological Longevity by Dietary Restriction in Budding Yeast’. Aging Cell 17(3). doi:10.1111/ACEL.12749.

11. Caydasi, Ayse Koca, Anton Khmelinskii, Zoulfia Darieva, Bahtiyar Kurtulmus, Michael Knop, and Gislene Pereira. 2023. ‘SWR1 Chromatin Remodeling Complex Prevents Mitotic Slippage during Spindle Position Checkpoint Arrest’. Molecular Biology of the Cell 34(2). doi:10.1091/MBC.E20-03-0179.

12. Creighton, Samantha D., Gilda Stefanelli, Mark A. Brimble, Luca Hategan, Emily Collins, Jacqueline Zakaria, Stefan Vislavski, Anas Reda, Timothy AB McLean, Brandon J. Walters, Margaret Fahnestock, Elias Orouji, and Iva Zovkic. 2023. ‘Sex-specific Accumulation and Therapeutic Effect of the Histone Variant H2A.Z in Alzheimer’s Disease’. Alzheimer’s & Dementia 19(S1). doi:10.1002/alz.064832.

13. Cruz-Bonilla, Erika, Sergio E. Campos, Soledad Funes, Cei Abreu-Goodger, and Alexander DeLuna. 2025. ‘The Core Genetic Drivers of Chronological Aging in Yeast Are Universal Regulators of Longevity’. Microbial Cell 12:274–89. doi:10.15698/MIC2025.10.861.

14. Dang, Weiwei, Kristan K. Steffen, Rocco Perry, Jean A. Dorsey, F. Brad Johnson, Ali Shilatifard, Matt Kaeberlein, Brian K. Kennedy, and Shelley L. Berger. 2009. ‘Histone H4 Lysine 16 Acetylation Regulates Cellular Lifespan’. Nature 459(7248):802–7. doi:10.1038/NATURE08085.

15. Dixon, Jesse R., Inkyung Jung, Siddarth Selvaraj, Yin Shen, Jessica E. Antosiewicz-Bourget, Ah Young Lee, Zhen Ye, Audrey Kim, Nisha Rajagopal, Wei Xie, Yarui Diao, Jing Liang, Huimin Zhao, Victor V. Lobanenkov, Joseph R. Ecker, James A. Thomson, and Bing Ren. 2015. ‘Chromatin Architecture Reorganization during Stem Cell Differentiation’. Nature 518(7539):331–36. doi:10.1038/NATURE14222.

16. Fabrizio, Paola, Shawn Hoon, Mehrnaz Shamalnasab, Abdulaye Galbani, Min Wei, Guri Giaever, Corey Nislow, and Valter D. Longo. 2010. ‘Genome-Wide Screen in Saccharomyces Cerevisiae Identifies Vacuolar Protein Sorting, Autophagy, Biosynthetic, and TRNA Methylation Genes Involved in Life Span Regulation’. PLoS Genetics 6(7):1–14. doi:10.1371/JOURNAL.PGEN.1001024.

17. Feser, Jason, David Truong, Chandrima Das, Joshua J. Carson, Jeffrey Kieft, Troy Harkness, and Jessica K. Tyler. 2010. ‘Elevated Histone Expression Promotes Life Span Extension’. Molecular Cell 39(5):724–35. doi:10.1016/j.molcel.2010.08.015.

18. Garay, Erika, Sergio E. Campos, Jorge González de la Cruz, Ana P. Gaspar, Adrian Jinich, and Alexander DeLuna. 2014. ‘High-Resolution Profiling of Stationary-Phase Survival Reveals Yeast Longevity Factors and Their Genetic Interactions’. PLoS Genetics 10(2). doi:10.1371/JOURNAL.PGEN.1004168.

19. Gerashchenko, Maxim V., Zalan Peterfi, Sun Hee Yim, and Vadim N. Gladyshev. 2021. ‘Translation Elongation Rate Varies among Organs and Decreases with Age’. Nucleic Acids Research 49(2):E9. doi:10.1093/NAR/GKAA1103.

20. Giaimo, Benedetto Daniele, Francesca Ferrante, Andreas Herchenröther, Sandra B. Hake, and Tilman Borggrefe. 2019. ‘The Histone Variant H2A.Z in Gene Regulation’. Epigenetics & Chromatin 12(1). doi:10.1186/S13072-019-0274-9.

21. Gietz, R. Daniel. 2014. ‘Yeast Transformation by the LiAc/SS Carrier DNA/PEG Method’. Methods in Molecular Biology (Clifton, N.J.) 1205. doi:10.1007/978-1-4939-1363-3_1.

22. Gingold, Hila, and Yitzhak Pilpel. 2011. ‘Determinants of Translation Efficiency and Accuracy’. Molecular Systems Biology 7. doi:10.1038/MSB.2011.14.

23. Gonskikh, Yulia, and Norbert Polacek. 2017. ‘Alterations of the Translation Apparatus during Aging and Stress Response’. Mechanisms of Ageing and Development 168:30–36. doi:10.1016/J.MAD.2017.04.003.

24. Gresham, David, Viktor M. Boer, Amy Caudy, Naomi Ziv, Nathan J. Brandt, John D. Storey, and David Botstein. 2011. ‘System-Level Analysis of Genes and Functions Affecting Survival during Nutrient Starvation in Saccharomyces Cerevisiae’. Genetics 187(1):299–317. doi:10.1534/GENETICS.110.120766.

25. Hansen, Malene, Stefan Taubert, Douglas Crawford, Nataliya Libina, Seung Jae Lee, and Cynthia Kenyon. 2007. ‘Lifespan Extension by Conditions That Inhibit Translation in Caenorhabditis Elegans’. Aging Cell 6(1):95–110. doi:10.1111/J.1474-9726.2006.00267.X.

26. Hong, Jingjun, Hanqiao Feng, Feng Wang, Anand Ranjan, Jianhong Chen, Jiansheng Jiang, Rodolfo Ghirlando, T. Sam Xiao, Carl Wu, and Yawen Bai. 2014. ‘The Catalytic Subunit of the SWR1 Remodeler Is a Histone Chaperone for the H2A.Z-H2B Dimer’. Molecular Cell 53(3):498–505. doi:10.1016/J.MOLCEL.2014.01.010.

27. Howard, Amber C., Jarod Rollins, Santina Snow, Sarah Castor, and Aric N. Rogers. 2016. ‘Reducing Translation through EIF4G/IFG-1 Improves Survival under ER Stress That Depends on Heat Shock Factor HSF-1 in Caenorhabditis Elegans’. Aging Cell 15(6):1027–38. doi:10.1111/ACEL.12516.

28. Hsu, Chih Chao, Jiejun Shi, Chao Yuan, Dan Zhao, Shiming Jiang, Jie Lyu, Xiaolu Wang, Haitao Li, Hong Wen, Wei Li, and Xiaobing Shi. 2018. ‘Recognition of Histone Acetylation by the GAS41 YEATS Domain Promotes H2A.Z Deposition in Non-Small Cell Lung Cancer’. Genes and Development 32(1):58–69. doi:10.1101/GAD.303784.117/-/DC1.

29. Hu, Zheng, Bo Xia, Spike D. L. Postnikoff, Zih Jie Shen, Alin S. Tomoiaga, Troy A. Harkness, Ja Hwan Seol, Wei Li, Kaifu Chen, and Jessica K. Tyler. 2018. ‘Ssd1 and Gcn2 Suppress Global Translation Efficiency in Replicatively Aged Yeast While Their Activation Extends Lifespan’. ELife 7. doi:10.7554/ELIFE.35551.

30. Jonak, Katarzyna, Ida Suppanz, Julian Bender, Agnieszka Chacinska, Bettina Warscheid, and Ulrike Topf. 2024. ‘Ageing-Dependent Thiol Oxidation Reveals Early Oxidation of Proteins with Core Proteostasis Functions’. Life Science Alliance 7(5). doi:10.26508/LSA.202302300.

31. Kmiecik, Szymon W., and Matthias P. Mayer. 2022. ‘Molecular Mechanisms of Heat Shock Factor 1 Regulation’. Trends in Biochemical Sciences 47(3):218–34. doi:10.1016/j.tibs.2021.10.004.

32. Kobor, Michael S., Shivkumar Venkatasubrahmanyam, Marc D. Meneghini, Jennifer W. Gin, Jennifer L. Jennings, Andrew J. Link, Hiten D. Madhani, and Jasper Rine. 2004. ‘A Protein Complex Containing the Conserved Swi2/Snf2-Related ATPase Swr1p Deposits Histone Variant H2A.Z into Euchromatin’. PLoS Biology 2(5). doi:10.1371/JOURNAL.PBIO.0020131.

33. Komar, Anton A. 2009. ‘A Pause for Thought along the Co-Translational Folding Pathway’. Trends in Biochemical Sciences 34(1):16–24. doi:10.1016/J.TIBS.2008.10.002.

34. Koplin, Ansgar, Steffen Preissler, Yulia Llina, Miriam Koch, Annika Scior, Marc Erhardt, and Elke Deuerling. 2010a. ‘A Dual Function for Chaperones SSB–RAC and the NAC Nascent Polypeptide–Associated Complex on Ribosomes’. The Journal of Cell Biology 189(1):57. doi:10.1083/JCB.200910074.

35. Koplin, Ansgar, Steffen Preissler, Yulia Llina, Miriam Koch, Annika Scior, Marc Erhardt, and Elke Deuerling. 2010b. ‘A Dual Function for Chaperones SSB–RAC and the NAC Nascent Polypeptide–Associated Complex on Ribosomes’. The Journal of Cell Biology 189(1):57. doi:10.1083/JCB.200910074.

36. Lin, Chia Liang, Yuriy Chaban, David M. Rees, Elizabeth A. McCormack, Lorraine Ocloo, and Dale B. Wigley. 2017. ‘Functional Characterization and Architecture of Recombinant Yeast SWR1 Histone Exchange Complex’. Nucleic Acids Research 45(12):7249–60. doi:10.1093/NAR/GKX414.

37. Liti, Gianni, David M. Carter, Alan M. Moses, Jonas Warringer, Leopold Parts, Stephen A. James, Robert P. Davey, Ian N. Roberts, Austin Burt, Vassiliki Koufopanou, Isheng J. Tsai, Casey M. Bergman, Douda Bensasson, Michael J. T. O’Kelly, Alexander Van Oudenaarden, David B. H. Barton, Elizabeth Bailes, Alex N. Nguyen, Matthew Jones, Michael A. Quail, Ian Goodhead, Sarah Sims, Frances Smith, Anders Blomberg, Richard Durbin, and Edward J. Louis. 2009. ‘Population Genomics of Domestic and Wild Yeasts’. Nature 458(7236):337. doi:10.1038/NATURE07743.

38. Livak, Kenneth J., and Thomas D. Schmittgen. 2001. ‘Analysis of Relative Gene Expression Data Using Real-Time Quantitative PCR and the 2(-Delta Delta C(T)) Method’. Methods (San Diego, Calif.) 25(4):402–8. doi:10.1006/METH.2001.1262.

39. Longo, Valter D., and Paola Fabrizio. 2012. ‘Chronological Aging in Saccharomyces Cerevisiae’. Sub-Cellular Biochemistry 57:101–21. doi:10.1007/978-94-007-2561-4_5.

40. López-Otín, Carlos, Maria A. Blasco, Linda Partridge, Manuel Serrano, and Guido Kroemer. 2013. ‘The Hallmarks of Aging’. Cell 153(6):1194. doi:10.1016/J.CELL.2013.05.039.

41. Mahapatra, Sahasransu, Pooran S. Dewari, Anubhav Bhardwaj, and Purnima Bhargava. 2011. ‘Yeast H2A.Z, FACT Complex and RSC Regulate Transcription of TRNA Gene through Differential Dynamics of Flanking Nucleosomes’. Nucleic Acids Research 39(10):4023–34. doi:10.1093/NAR/GKQ1286.

42. Malik, Yasir, Yavuz Kulaberoglu, Shajahan Anver, Sara Javidnia, Gillian Borland, Rene Rivera, Stephen Cranwell, Danel Medelbekova, Tatiana Svermova, Jackie Thomson, Susan Broughton, Tobias von der Haar, Colin Selman, Jennifer M. A. Tullet, and Nazif Alic. 2024. ‘Disruption of TRNA Biogenesis Enhances Proteostatic Resilience, Improves Later-Life Health, and Promotes Longevity’. PLoS Biology 22(10). doi:10.1371/JOURNAL.PBIO.3002853.

43. Matecic, Mirela, Daniel L. Smith, Xuewen Pan, Nazif Maqani, Stefan Bekiranov, Jef D. Boeke, and Jeffrey S. Smith. 2010. ‘A Microarray-Based Genetic Screen for Yeast Chronological Aging Factors’. PLoS Genetics 6(4). doi:10.1371/JOURNAL.PGEN.1000921.

44. Mizuguchi, Gaku, Xuetong Shen, Joe Landry, Wei Hua Wu, Subhojit Sen, and Carl Wu. 2004. ‘ATP-Driven Exchange of Histone H2AZ Variant Catalyzed by SWR1 Chromatin Remodeling Complex’. Science (New York, N.Y.) 303(5656):343–48. doi:10.1126/SCIENCE.1090701.

45. Morillo-Huesca, Macarena, Marta Clemente-Ruiz, Eloísa Andújar, and Félix Prado. 2010. ‘The SWR1 Histone Replacement Complex Causes Genetic Instability and Genome-Wide Transcription Misregulation in the Absence of H2A.Z’. PloS One 5(8). doi:10.1371/JOURNAL.PONE.0012143.

46. Papamichos-Chronakis, Manolis, Jocelyn E. Krebs, and Craig L. Peterson. 2006. ‘Interplay between Ino80 and Swr1 Chromatin Remodeling Enzymes Regulates Cell Cycle Checkpoint Adaptationin Response to DNA Damage’. Genes & Development 20(17):2437. doi:10.1101/GAD.1440206.

47. Parker, Steven, Marcin G. Fraczek, Jian Wu, Sara Shamsah, Alkisti Manousaki, Kobchai Dungrattanalert, Rogerio Alves De Almeida, Diego Estrada-Rivadeneyra, Walid Omara, Daniela Delneri, and Raymond T. O’keefe. 2017. ‘A Resource for Functional Profiling of Noncoding RNA in the Yeast Saccharomyces Cerevisiae’. RNA (New York, N.Y.) 23(8):1166–71. doi:10.1261/RNA.061564.117.

48. Parker, Steven, Marcin G. Fraczek, Jian Wu, Sara Shamsah, Alkisti Manousaki, Kobchai Dungrattanalert, Rogerio Alves de Almeida, Edith Invernizzi, Tim Burgis, Walid Omara, Sam Griffiths-Jones, Daniela Delneri, and Raymond T. O’Keefe. 2018a. ‘Large-Scale Profiling of Noncoding RNA Function in Yeast’. PLoS Genetics 14(3). doi:10.1371/JOURNAL.PGEN.1007253.

49. Parker, Steven, Marcin G. Fraczek, Jian Wu, Sara Shamsah, Alkisti Manousaki, Kobchai Dungrattanalert, Rogerio Alves de Almeida, Edith Invernizzi, Tim Burgis, Walid Omara, Sam Griffiths-Jones, Daniela Delneri, and Raymond T. O’Keefe. 2018b. ‘Large-Scale Profiling of Noncoding RNA Function in Yeast’. PLoS Genetics 14(3). doi:10.1371/JOURNAL.PGEN.1007253.

50. Pegoraro, Gianluca, and Tom Misteli. 2009. ‘The Central Role of Chromatin Maintenance in Aging’. Aging 1(12):1017–22. doi:10.18632/AGING.100106.

51. Percudani, Riccardo, Angelo Pavesi, and Simone Ottonello. 1997. ‘Transfer RNA Gene Redundancy and Translational Selection in Saccharomyces Cerevisiae’. Journal of Molecular Biology 268(2):322–30. doi:10.1006/JMBI.1997.0942.

52. Romila, Catalina A., St John Townsend, Michal Malecki, Stephan Kamrad, María Rodríguez- López, Olivia Hillson, Cristina Cotobal, Markus Ralser, and Jürg Bähler. 2021. ‘Barcode Sequencing and a High-Throughput Assay for Chronological Lifespan Uncover Ageing-Associated Genes in Fission Yeast’. Microbial Cell 8(7):146–60. doi:10.15698/MIC2021.07.754.

53. Sagi, Dror, Roni Rak, Hila Gingold, Idan Adir, Gadi Maayan, Orna Dahan, Limor Broday, Yitzhak Pilpel, and Oded Rechavi. 2016. ‘Tissue– and Time-Specific Expression of Otherwise Identical TRNA Genes’. PLoS Genetics 12(8). doi:10.1371/JOURNAL.PGEN.1006264.

54. Schneider-Poetsch, Tilman, Jianhua Ju, Daniel E. Eyler, Yongjun Dang, Shridhar Bhat, William C. Merrick, Rachel Green, Ben Shen, and Jun O. Liu. 2010. ‘Inhibition of Eukaryotic Translation Elongation by Cycloheximide and Lactimidomycin’. Nature Chemical Biology 6(3):209–17. doi:10.1038/NCHEMBIO.304.

55. Stefanelli, Gilda, Amber B. Azam, Brandon J. Walters, Mark A. Brimble, Caroline P. Gettens, Pascale Bouchard-Cannon, Hai Ying M. Cheng, Andrew M. Davidoff, Klotilda Narkaj, Jeremy J. Day, Andrew J. Kennedy, and Iva B. Zovkic. 2018. ‘Learning and Age-Related Changes in Genome-Wide H2A.Z Binding in the Mouse Hippocampus’. Cell Reports 22(5):1124–31. doi:10.1016/J.CELREP.2018.01.020.

56. Stein, Kevin C., Fabián Morales-Polanco, Joris van der Lienden, T. Kelly Rainbolt, and Judith Frydman. 2022. ‘Ageing Exacerbates Ribosome Pausing to Disrupt Cotranslational Proteostasis’. Nature 601(7894):637–42. doi:10.1038/S41586-021-04295-4.

57. Svotelis, Amy, Nicolas Gévry, Gilles Grondin, and Luc Gaudreau. 2010. ‘H2A.Z Overexpression Promotes Cellular Proliferation of Breast Cancer Cells’. Cell Cycle 9(2):364–70. doi:10.4161/CC.9.2.10465.

58. Swer, Pynskhem Bok, and Ramesh Sharma. 2021. ‘ATP-Dependent Chromatin Remodelers in Ageing and Age-Related Disorders’. Biogerontology 22(1). doi:10.1007/S10522-020-09899-3.

59. Tong, Amy Hin Yan, and Charles Boone. 2006. ‘Synthetic Genetic Array Analysis in Saccharomyces Cerevisiae’. Methods in Molecular Biology (Clifton, N.J.) 313:171–92. doi:10.1385/1-59259-958-3:171.

60. Torrent, Marc, Guilhem Chalancon, Natalia S. De Groot, Arthur Wuster, and M. Madan Babu. 2018. ‘Cells Alter Their TRNA Abundance to Selectively Regulate Protein Synthesis during Stress Conditions’. Science Signaling 11(546). doi:10.1126/SCISIGNAL.AAT6409.

61. Tuller, Tamir, Asaf Carmi, Kalin Vestsigian, Sivan Navon, Yuval Dorfan, John Zaborske, Tao Pan, Orna Dahan, Itay Furman, and Yitzhak Pilpel. 2010. ‘An Evolutionarily Conserved Mechanism for Controlling the Efficiency of Protein Translation’. Cell 141(2):344–54. doi:10.1016/J.CELL.2010.03.031.

62. Tyczewska, Agata, and Kamilla Grzywacz. 2023. ‘TRNA-Derived Fragments as New Players in Regulatory Processes in Yeast’. Yeast 40(8):283–89. doi:10.1002/YEA.3829.

63. Tye, Blake W., and L. Stirling Churchman. 2021. ‘Hsf1 Activation by Proteotoxic Stress Requires Concurrent Protein Synthesis’. Molecular Biology of the Cell 32(19):1800–1806. doi:10.1091/MBC.E21-01-0014.

64. Wu, Wei Hua, Samar Alami, Edward Luk, Chwen Huey Wu, Subhojit Sen, Gaku Mizuguchi, Debbie Wei, and Carl Wu. 2005. ‘Swc2 Is a Widely Conserved H2AZ-Binding Module Essential for ATP-Dependent Histone Exchange’. Nature Structural & Molecular Biology 12(12):1064–71. doi:10.1038/NSMB1023.

65. Xu, Bowen, Alexander Hull, Olivia N. M. Hill, Naja Kobal, Enric Ureña, Linda Partridge, and Nazif Alic. 2025. ‘Loss of Pol III Repressor Maf1 in Neurons Promotes Longevity by Preventing the Age-Related Decline in 5S RRNA and Translation’. PLoS Biology 23(7). doi:10.1371/JOURNAL.PBIO.3003250.

66. Yen, Kuangyu, Vinesh Vinayachandran, and B. Franklin Pugh. 2013. ‘XSWR-C and INO80 Chromatin Remodelers Recognize Nucleosome-Free Regions near +1 Nucleosomes’. Cell 154(6):1246. doi:10.1016/j.cell.2013.08.043.

67. Zhang, Gong, and Zoya Ignatova. 2011. ‘Folding at the Birth of the Nascent Chain: Coordinating Translation with Co-Translational Folding’. Current Opinion in Structural Biology 21(1):25–31. doi:10.1016/j.sbi.2010.10.008.

68. Zhou, Bo O., Shan-Shan Wang, Lu-Xia Xu, Fei-Long Meng, Yao-Ji Xuan, Yi-Min Duan, Jian-Yong Wang, Hao Hu, Xianchi Dong, Jianping Ding, and Jin-Qiu Zhou. 2010. ‘SWR1 Complex Poises Heterochromatin Boundaries for Antisilencing Activity Propagation’. Molecular and Cellular Biology 30(10):2391. doi:10.1128/MCB.01106-09.

69. Zhou, Zheng, Bao Sun, Dongsheng Yu, and Meng Bian. 2021. ‘Roles of TRNA Metabolism in Aging and Lifespan’. Cell Death & Disease 12(6). doi:10.1038/S41419-021-03838-X.

