## Supporting Information for "SWR1C loss promotes longevity through tRNA-mediated proteostasis"

**Figure S1.** Absence of *SWR1* extends CLS in different genetic backgrounds of *S. cerevisiae*

**Figure S2.** SGA-based construction of gene-deletion strain arrays for competitive-CLS profiling

**Figure S3.** Competitive-CLS screening yields highly reproducible CLS measurements

**Figure S4.** Epistasis analysis of *SWR1*-impaired strains and selected tRNA mutants

**Table S1.** GO terms enriched in *SWR1*-epistasis analysis

**Table S2.** Media Recipes

**Table S3.** PCR primers used for tRNA quantification

**Dataset S1.** CLS-epistasis screening of protein-coding genes

**Dataset S2.** CLS screening of ncRNA genes

**Dataset S3.** CLS-epistasis screening of ncRNA genes

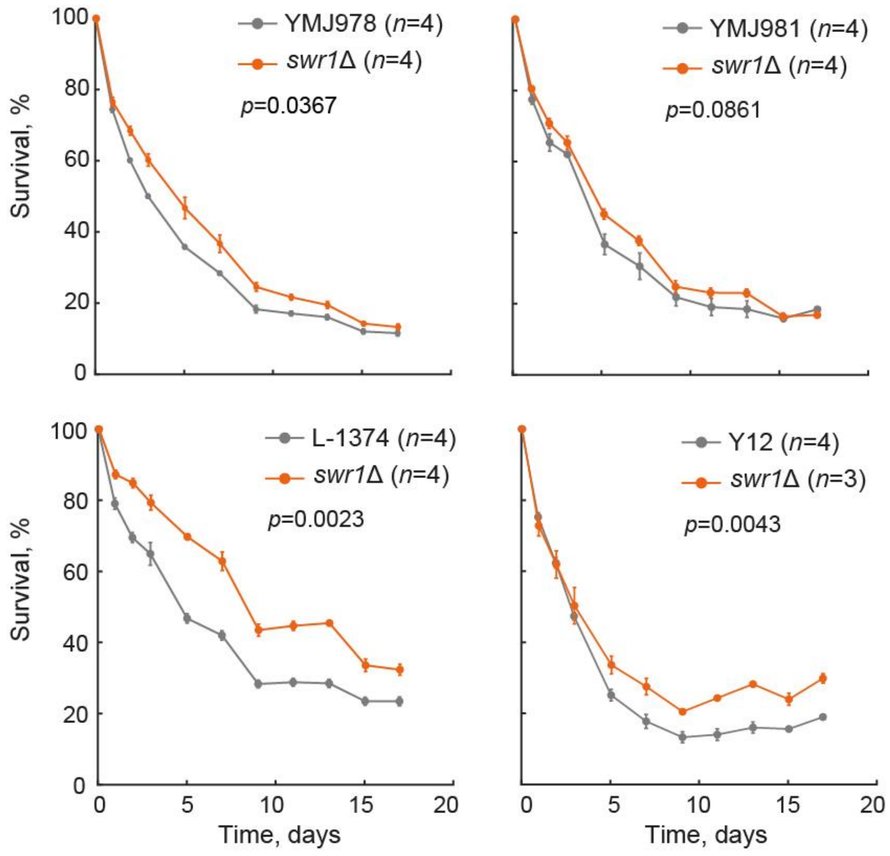


**Figure S1. Absence of SWR1 extends CLS in different genetic backgrounds of *S. cerevisiae.*** Survival curves were obtained by monitoring outgrowth kinetics at multiple time points from stationary-phase cultures (Murakami and Kaeberlein, 2009). For each wild parental strain (Liti *et al.*, 2009) deletions of *HO* (neutral insertion) and *SWR1* were generated via PCR-directed mutagenesis. Three to four independent colonies were isolated for each transformation and cultures were subsequently aged in 800 µL of SC aging medium at 30 °C without shaking. At each sampling point starting 24 hours after inoculation (*Time*=0 d), 15 µL of aging culture was transferred to 96-well plates containing 135 µL of fresh YNB-lf medium. Optical density ($OD_{600}$) was measured in a multilabel plate reader integrated to an automated robotic system. To estimate survival relative to $T$=0, outgrowth kinetics were recorded hourly until saturation. An AUC was calculated for each strain replicate; statistical significance between *hoΔ* and *swr1Δ* strains was determined using a two-tailed Student's *t*-test.

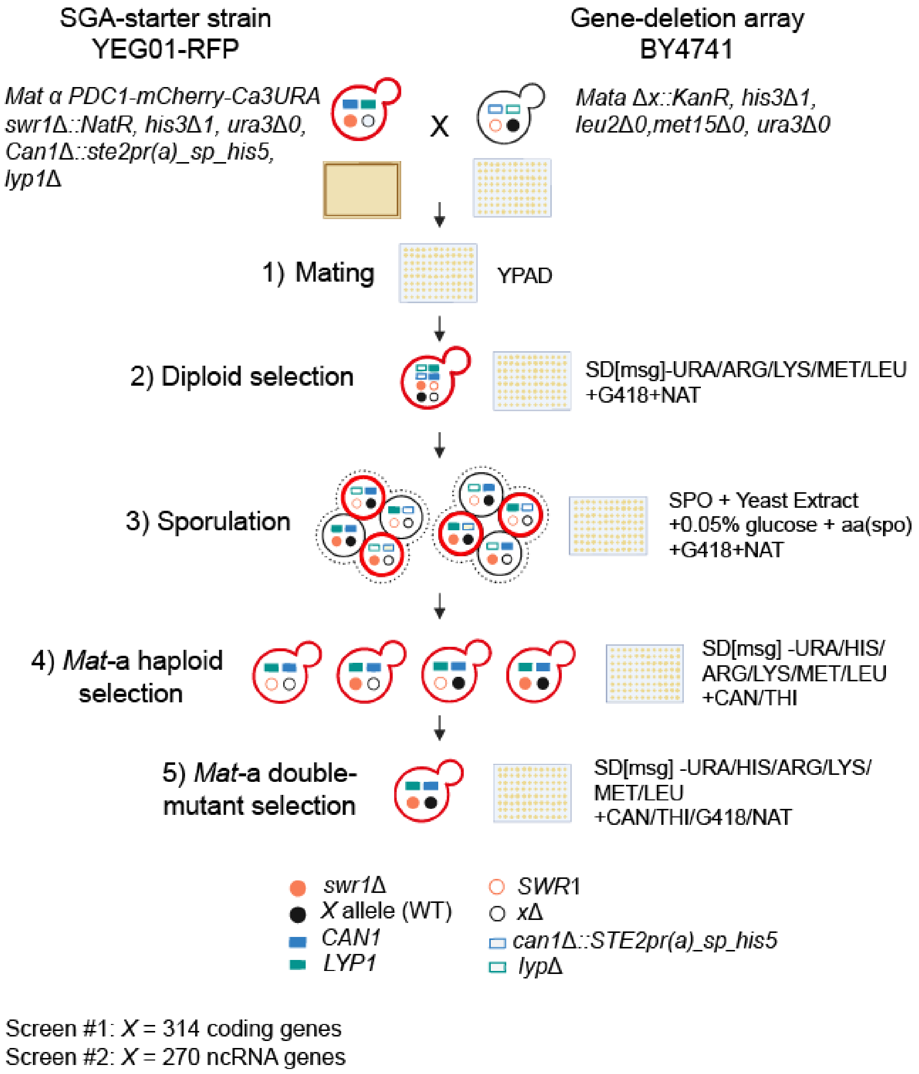


**Figure S2. SGA-based construction of gene-deletion strain arrays for competitive-CLS profiling.** *MATα* query strains carried an mCherry tag (RFP) and either an *HO* neutral insertion for single deletions or an *SWR1* insertion for double deletions (orange circles, shown here) each linked to the *NatMX4* cassette, and were crossed to arrays of 314 *MATa* coding-gene deletions or 270 *MATa* non-coding RNA deletions, each linked to *KanMX4* (black circles). The starter YEG01-RFP strain also had two additional markers: *can1Δ::STE2pr-Sp_his5* (blue rectangle) and *lyp1Δ* (green rectangle), used for selection of haploid progeny. After mating and diploid selection, sporulation was carried out for ten days at 21 °C. Double-mutant RFP haploids were selected on media with G418 and ClonNAT, lacking uracil and histidine, and supplemented with canavanine (CAN) and thialysine (THI) to counterselect for diploids. For competitive-CLS screening, *hoΔ* or *swr1Δ* references were generated starting with the YEG02-CFP crossed to a *his3Δ* neutral-insertion strain. Method and figure adapted from (Tong and Boone, 2006).

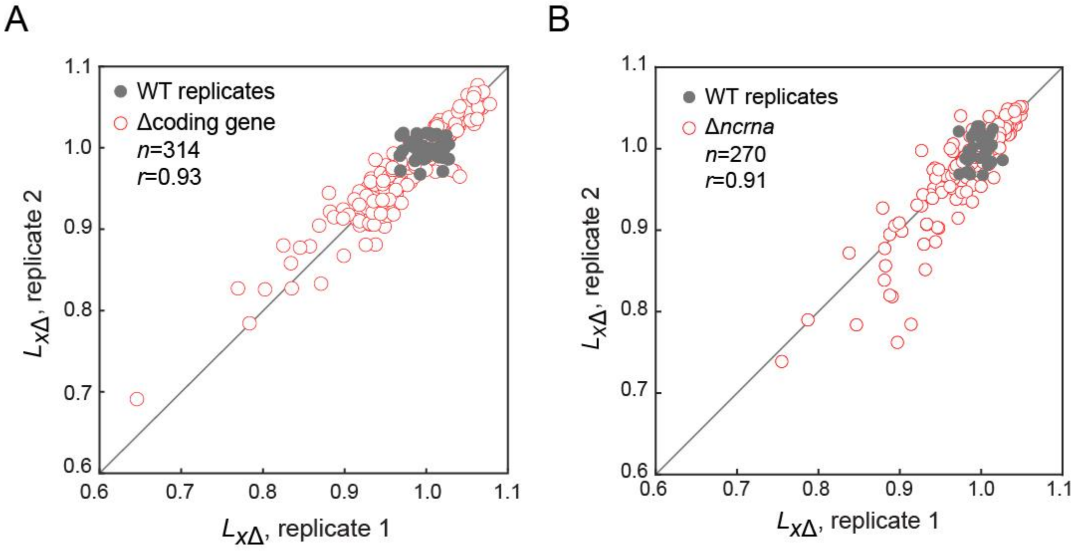


**Figure S3. Competitive-CLS screening yields highly reproducible CLS measurements.** Scatter plots show the correlation coefficient (*r*) between CLS values ($L_{x\Delta}$) obtained from two independent competitive-aging assays. Closed gray circles in both panels represent CLS ($L_{x\Delta}$) values for WT replicates. **(*A*)** Correlation of CLS ($L_{x\Delta}$) values for 314 deletion strains of coding genes (open red circles). **(*B*)** Correlation of CLS ($L_{x\Delta}$) values for 270 deletion strains of non-coding RNA genes (open red circles).

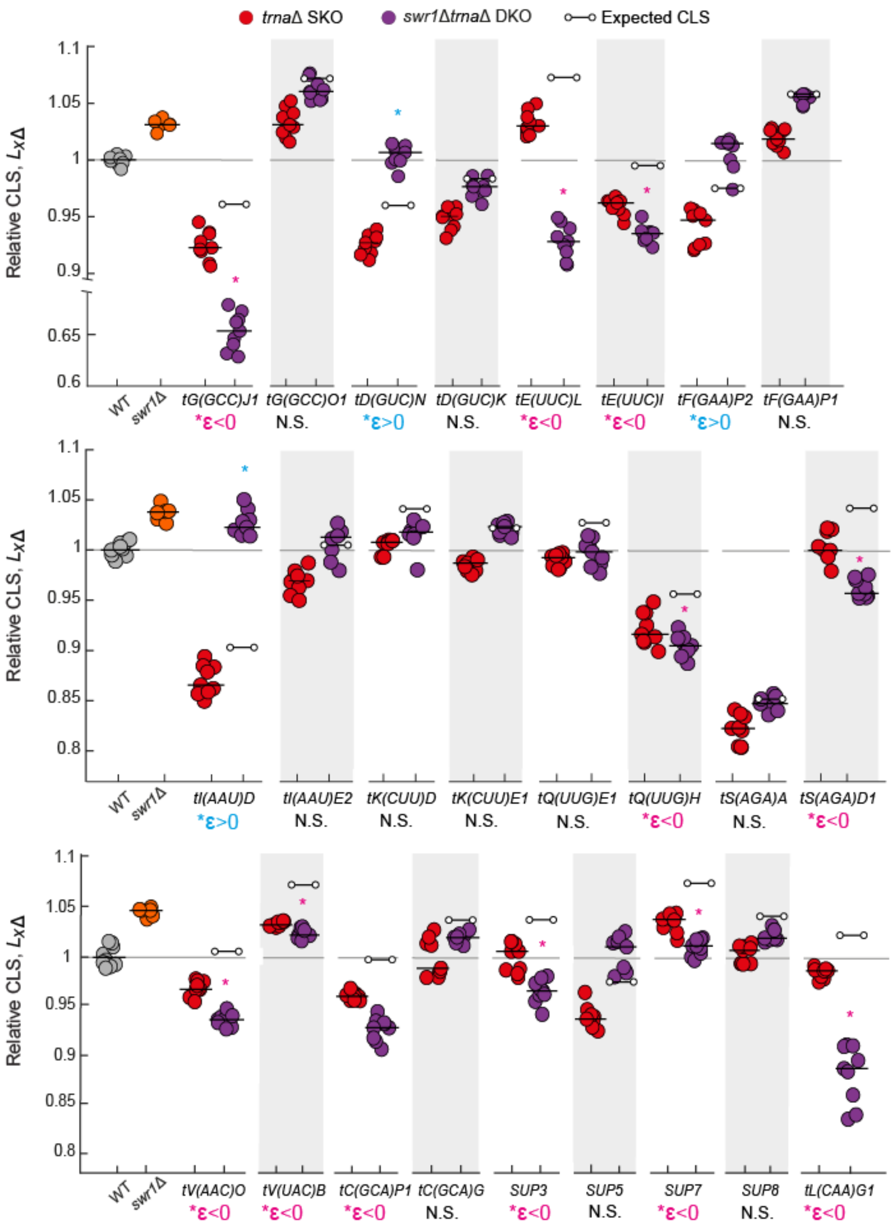


**Figure S4. Epistasis analysis of *SWR1*-impaired strains and selected tRNA mutants.** Plots show the measured relative lifespan (CLS) of selected gene pairs (single and double knockouts). Black lines represent the median of 9 replicates per gene-pair strain, and 6-10 replicates for WT and *swr1Δ* controls. Each circle-endpoint line (o—o) indicates the expected CLS for the $L_{x\Delta,swr1\Delta}$ double mutant, based on the multiplicative value of the corresponding single mutants $(L_{x\Delta})\cdot(L_{swr1\Delta})$. Significant positive and negative epistatic interactions were determined by two-tailed *t*-tests (*$p$<0.05). Each plot is an experimental batch sharing the same WT and *swr1Δ* strains.

**Table S1. GO terms enriched in *SWR1*-epistasis analysis**

| GO term [1] | Description | Genes | *p-val* [2] | *n* [3] |
|---|---|---|---|---|
| GO:0000122 | Negative regulation of transcription by RNA polymerase II | *BMH1, CUP9, DEP1, DIG2, FKH2, GAL80, GAT1, GIS1, GLN3, HDA2, ISW2, OPI1, PHO85, RXT2, UME6, WTM2* | 0.037 | 16 |
| GO:0006364 | rRNA processing | *RPS14A, RPS21A, RPS6B, RRP8, SSF1* | 0.031 | 5 |
| GO:0006417 | Translation | *RPS14A, RPS21A, RPS6B, RRP8, RPS18A, RPS1A, TRM7* | 0.041 | 7 |
| GO:0007049 | Cell cycle | *BUD14, EGT2, FAR10, FAR11, FAR7, MEC3, PCL1, SUN4, TOR1, UTH1, VHS2* | 0.049 | 11 |
| GO:0031509 | Subtelomeric heterochromatin formation | *BR2, ISW2, MEC3, PNC1, RAD6, SPP1, SWD1* | 0.046 | 7 |
| GO:0034599 | Cellular response to oxidative stress | *MXR2, OCA1, PIM1, SCO1,TOR1,TRX3* | 0.037 | 6 |
| GO:0043161 | Proteasome-mediated ubiquitin-dependent protein catabolic process | *DOA1, PEX10, PRE9, RAD6, RMD5, UBC8, UBP14, UFD4* | 0.032 | 8 |
| GO:0071555 | Cell wall organization | *CCW12, CHS7, DSE2, SUN4, TOS1, UTH1* | 0.040 | 6 |
| GO:1902600 | Proton transmembrane transport | *ATP4, COX5A, COX6, QCR7, QCR9* | 0.045 | 5 |

1. GO annotations were downloaded from the Yeast Genome Database (Nov 2024)

2. *p*-value of the Wilcoxon rank sum test, compared to the *S* distribution of 314 genes screened

3. Number of genes screened within the enriched GO term

**Table S2. Media Recipes**

---

**Aging medium (SC)**

Recipe
For 1 L of Aging medium, mix together:
6.7 g Yeast Nitrogen Base w/o Amino Acids (Difco™, 291940)
2 g Amino acid complete mix.
20 g D-(+)- Glucose (Sigma-aldrich, G2870)
Add the components to 500 mL of deionized water and dissolve
Add deionized water to make 1 L of medium.
Filter sterilize

*We previously added 1.79 g of Uracil (Sigma-aldrich, U0750) the commercial Synthetic Drop-out mix (20 g) (Sigma-aldrich, Y1501) to obtain a complete amino acid mix
*Note: do not buffer pH*

---

**YPD**

Recipe
For 1 L of YPD medium, mix together:

10 g Yest extract (Difco™)
20 g Peptone (Difco™)
20 g D-(+)-Glucose
20 g Agar (Difco™) (for plates only)
Dissolve components in 1 L of deionized water distilled
Autoclave for 20 min at 121°C and 0.5 bar.

---

**SD-URA**

Recipe
2% glucose
6.7g Y.N.B. w/o amino acids
Dissolve components in 1 L of deionized water distilled
Autoclave for 20 min at 121°C and 0.5 bar.

Add
**Amino acids stock:**
20 mg/L histidine, 120 mg/L leucine, 60 mg/L lysine, 20 mg/L tryptophan, 20 mg/L methionine and 20 mg/L adenine

---

**Media for SGA**

**SGA-starter medium**

Recipe
SD [msg] -URA/ARG/LYS/MET/LEU + canavanine/thialysine/ClonNAT (1L)

1.7g Y.N.B. w/o amino acids and ammonium sulfate
1g Monosodium glutamic acid

1.6g Amino acid drop-out mix -URA /HIS/ARG/LYS/MET/LEU
2 mL His stock (10 mg·mL$^{-1}$)
Dissolve in 200 mL of $H_2O$ and filter sterilize
20g Agar
Dissolve in 700 mL of $H_2O$
After autoclaved, mix drop-out with agar, and add 100 mL 20% glucose, and 1 mL of canavanine solution (100 mg·mL$^{-1}$), 1 mL of thialysine solution (100 mg·mL$^{-1}$), 1 mL of ClonNAT solution (100 mg·mL$^{-1}$).

---

**Target array medium**

Recipe
YPAD + G418 + AMP (1L)
120mg Adenine Sulfate dehydrate
10g Yeast extract
20g Peptone
20g Agar
Dissolve in 900 mL of $H_2O$ in a 1L flask, add 100 mL of 20% glucose after autoclaving, 1 mL G418 solution (200 mg·mL$^{-1}$), and 1 mL Ampicillin solution (100 mg·mL$^{-1}$).

---

**Diploid selection medium**

Recipe
SD [msg] -URA /ARG/LYS/MET/LEU+G418+ClonNAT (1L)
1.7g Y.N.B. w/o amino acids and ammonium sulfate
1g Monosodium glutamic acid
1.6g Amino acid drop-out mix -URA /HIS/ARG/LYS/MET/LEU
2 mL His stock
Dissolve in 200 mL of $H_2O$ and filter sterilize
20g Agar
Dissolve in 700 mL of $H_2O$ after autoclaved, mix drop-out with agar, and add 100 mL 20% glucose, add 1 mL of G418 and 1 mL of ClonNAT

---

**Sporulation medium**

Recipe

SPO + Y.E. + glucose + amino acids + G418 (1L)
10g Potassium acetate
1g Yeast extract
0.5g Glucose
0.1g Amino acid supplement for SPO medium
20g Agar
Dissolve in 1L of deionized $H_2O$, and 250 µL of G418 solution

Amino-acid supplement for sporulation medium
2g Histidine
10g Leucine
2g Lysine
2g Uracil

---

Mix thoroughly by turning end-over-end for at least 15min with clean marble.
Store in tinted glass bottle at room temperature.

---

**MAT a haploid selection medium**

Recipe
SD-HIS/ARG/LYS/LEU/MET + canavanine/thialysine (1L)
6.7g Y.N.B. w/o amino acids
1.6g Amino acid drop-out mix -URA/HIS/ARG/LYS/LEU/MET
10 mL Ura stock (2 mg·$mL^{-1}$. pH 9.0)
Dissolve in 100 mL of $H_2O$ and filter sterilize
20g Agar
Dissolve in 800mL of $H_2O$ in a 2L flask.
After autoclaved, mix drop-out with agar, and add 100 mL 20% glucose, 1mL of canavanine solution, and 1 mL of thialysine solution.

---

**kanR selection medium**

Recipe
SD [msg]-HIS/ARG/LYS/LEU/MET + canavanine/thialysine/G418 (1L)
1.7g Y.N.B. w/o amino acids and ammonium sulfate
1g Monosodium glutamic acid
1.6g Amino acid drop-out mix -URA/HIS/ARG/LYS/LEU/MET
10 mL Ura stock
Dissolve in 200 mL of $H_2O$ and filter sterilize
20g Agar
Dissolve in 700 ml of $H_2O$
After autoclaved, mix drop-out with agar, and add 100 mL 20% glucose, 1 mL of canavanine solution, 1 mL of thialysine solution, and 1 mL of G418.

---

**kanR/NatR/Ca3URA selection medium**

Recipe
SD [msg]-HIS/ARG/LYS/LEU/MET/URA + canavanine/thialysine/G418/ClonNAT (1L)

1.7g Y.N.B. w/o amino acids and ammonium sulfate
1g Monosodium glutamic acid
1.6g Amino acid drop-out mix -URA/HIS/ARG/LYS/LEU/MET
Dissolve in 200mL of $H_2O$ and filter sterilize
20g Agar
Dissolve in 700 mL of $H_2O$
After autoclaved, mix drop-out with agar, and add 100 mL 20% glucose, and 1 mL of canavanine solution, 1 mL of thialysine solution, 1 mL of G418, and 1 mL ClonNAT

---

**Table S3. PCR primers used for tRNA quantification [1]**

| Amino acid | tRNA Anticodon | Forward 5'- 3' | Reverse 5'- 3' |
|---|---|---|---|
| Cys | GCA | CTCGTATGGCGCAGTGGTAG | CTCGCACTCAGGATCGAACT |
| Gly | GCC | GCGCAAGTGGTTTAGTGGTA | GCAAGCCCGGAATCGAAC |
| Leu | CAA | TTTGGCCGAGCGGTCTAAGG | TGCATCTTACGATACCTGAGCTT |
| Lys | CUU | CTTGTTGGCGCAATCGGTAG | GGGCTCGAACCCCTAACCTT |
| Ser | AGA | ACTTGGCCGAGTGGTTAAGG | ACAACTGCAGGACTCGAACC |
| Trp | CCA | GAAGCGGTGGCTCAATGGTA | CGGACAGGAATTGAACCTGC |
| Tyr | GUA | AGCCAAGTTGGTTTAAGGCG | AGCCAAGTTGGTTTAAGGCG |
| Val | UAC | TGGTCCAGTGGTTCAAGACG | TTCGAACTCGGGATCTTCGC |
| *ACT1** | N/A | TGGATTCTGAGGTTGCTGCT | AGATGGGAAGACAGCACGAG |

1. Primers are from Torrent et al., 2018.

*This study
